# Intraoperative Bioprinting of Human Adipose-derived Stem cells and Extra-cellular Matrix Induces Hair Follicle-Like Downgrowths and Adipose Tissue Formation during Full-thickness Craniomaxillofacial Skin Reconstruction

**DOI:** 10.1101/2023.10.03.560695

**Authors:** Youngnam Kang, Miji Yeo, Irem Deniz Derman, Dino J. Ravnic, Yogendra Pratap Singh, Mecit Altan Alioglu, Yang Wu, Jasson Makkar, Ryan R. Driskell, Ibrahim T. Ozbolat

## Abstract

Craniomaxillofacial (CMF) reconstruction is a challenging clinical dilemma. It often necessitates skin replacement in the form of autologous graft or flap surgery, which differ from one another based on hypodermal/dermal content. Unfortunately, both approaches are plagued by scarring, poor cosmesis, inadequate restoration of native anatomy and hair, alopecia, donor site morbidity, and potential for failure. Therefore, new reconstructive approaches are warranted, and tissue engineered skin represents an exciting alternative. In this study, we demonstrated the reconstruction of CMF full-thickness skin defects using intraoperative bioprinting (IOB), which enabled the repair of defects via direct bioprinting of multiple layers of skin on immunodeficient rats in a surgical setting. Using a newly formulated patient-sourced allogenic bioink consisting of both human adipose-derived extracellular matrix (adECM) and stem cells (ADSCs), skin loss was reconstructed by precise deposition of the hypodermal and dermal components under three different sets of animal studies. adECM, even at a very low concentration such as 2% or less, has shown to be bioprintable via droplet-based bioprinting and exhibited *de novo* adipogenic capabilities both *in vitro* and *in vivo*. Our findings demonstrate that the combinatorial delivery of adECM and ADSCs facilitated the reconstruction of three full-thickness skin defects, accomplishing near-complete wound closure within two weeks. More importantly, both hypodermal adipogenesis and downgrowth of hair follicle-like structures were achieved in this two-week time frame. Our approach illustrates the translational potential of using human-derived materials and IOB technologies for full-thickness skin loss.

## 1. INTRODUCTION

Tissue engineered skin substitutes are viable alternatives to conventional reconstructive surgery and are often used as an alternative to autologous grafts and flaps. Grafts can be differentiated from flaps based on the hypodermal to dermal ratio. Unfortunately, both approaches are suboptimal and rooted in antiquated principles. However, most current tissue engineering approaches mainly focus on dermal replacement and neglect the underlying hypodermis, for a multitude of reasons [1,2].

Anatomically, skin is made up of three major layers; epidermis, the outermost layer; dermis, the middle layer; and hypodermis, the deepest fat layer[3]. The epidermis includes mainly keratinocytes and melanocytes, which are not directly involved in adipogenesis, while the dermis (including sweat glands, hair follicles and sebaceous glands) and the hypodermis (made up of connective tissue and fat) are reported to be directly involved in adipogenesis [4,5]. Specifically, the hypodermis connects to the underlying fascia [6] and contains adipocytes, blood vessels, nerves, and stromal cells, which are all crucial to normal mechanical and thermoregulatory maintenance[6,7]. Additionally, the hypodermis is essential to wound healing by supporting keratinocyte and fibroblast proliferation [8,9]. Recent studies have demonstrated that adipocytes regulate wound repair by mediating fibroblast recruitment. Thus, reconstruction of this adipose-laden compartment is highly crucial for systemic metabolism, immune function, and skin physiology.

Adipogenesis also has a critical role in hair follicle cycling, particularly, in extending the anagen stage of hair growth [10,11]. During the late anagen to middle telogen stage of hair follicle cycles, adipocytes tend to emit bone morphogenetic protein-2 (BMP-2), which helps the resting state of hair follicle stem cells and facilitates hair growth [12]. Recently, there has been increasing number of studies showing that dermal adipocytes are highly associated with hair follicles [13–17]. For example, Kim et al. demonstrated that adipose-derive stem cells (ADSCs) secrete various growth factors, such as platelet-derived growth factor (PDGF), vascular endothelial growth factor (VEGF), hepatocyte growth factor (HGF), keratinocyte growth factor (KGF) and fibroblast growth factor-1 and −2 (FGF-1and FGF-2), that have been shown to promote hair growth[18–22]. Moreover, the conditioned media of ADSCs induces hair growth in mice and human hair organ culture ex vivo [23]. ADSCs and their constituent extracts have also been tested in clinical trials for hair growth and has shown potential for hair regeneration [24–26].

The craniomaxillofacial (CMF) skeleton is particularly impacted by full thickness skin loss for both functional and cosmetic concerns. Skin is critical for the protection of vital structures, such as the brain and aerodigestive tract. Furthermore, any abnormality is conspicuous and can hamper normal social interactions. This makes CMF reconstruction following tumor extirpation or congenital deficiency particularly complex[27]. Despite many surgical advances, both autologous grafts and flaps continue to be plagued by scarring, poor cosmesis, inadequate restoration of native anatomy, alopecia, donor site morbidity, and potential for failure[1]. Therefore, new reconstructive approaches are warranted, and tissue engineered skin represents an exciting alternative. Tissue constructs can be made of acellular materials or synthesized from autologous, allograft, xenogeneic, or synthetic sources [28,29]. However, to date, a complete and competent full-thickness skin substitute is not available and that needs to be addressed prior to widespread clinical adoption.

In this study, we demonstrated intraoperative bioprinting (IOB) for the reconstruction of full-thickness CMF skin loss, where bioprinting was performed directly on the defect site, which is greatly beneficial as IOB facilitates the development of skin tissue heterogeneity in an anatomically accurate and cosmetically appropriate manner [30–32]. IOB has the capability to alleviate the need for invasive operations for fabrication of highly thin stratified structures and their transplantation experienced during conventional methods, such as employing wound dressings and injectable hydrogels. Indeed, it is extremely difficult to manually fabricate a stratified multilayer structure with 200-300 µm layer thickness using these conventional methods. Moreoever, IOB eliminates the need for shaping prefabricated wound dressings during the surgery and the precise stratified arrangement achieved during IOB also enables the successful crosslinking of deposited biomaterials *in situ*.

Using a micro-solenoid valve system, droplets of bioink solutions were ejected with precise control of size and reproducibility; and skin compartments were recreated using a newly formulated bioink with different concentrations of clinically-obtained human adipose-derived extracellular matrix (adECM) and human ADSCs on demand as shown in **Figure 1A**. In our study, only bilayered (dermis and hypodermis) constructs were deposited as it was expected that the epithelial cells could migrate on the bioprinted constructs and reconstitute the epidermal layer. To demonstrate the role of adECM and ADSCs for full-thickness skin reconstruction, we performed three different sets of animal studies as highlighted in **Figure 1B**. Our results indicate that intraoperatively bioprinted skin constructs enabled the reconstruction of full-thickness CMF skin defects. More importantly, the combinatorial delivery of adECM and ADSC in the hypodermis supported the formation of hypodermal adipose and promoted the development of hair follicle-like downgrowths, which will have a considerable impact on the translation of IOB of human surgical discards into clinic for repair of skin defects in the future.

**Figure 1.**
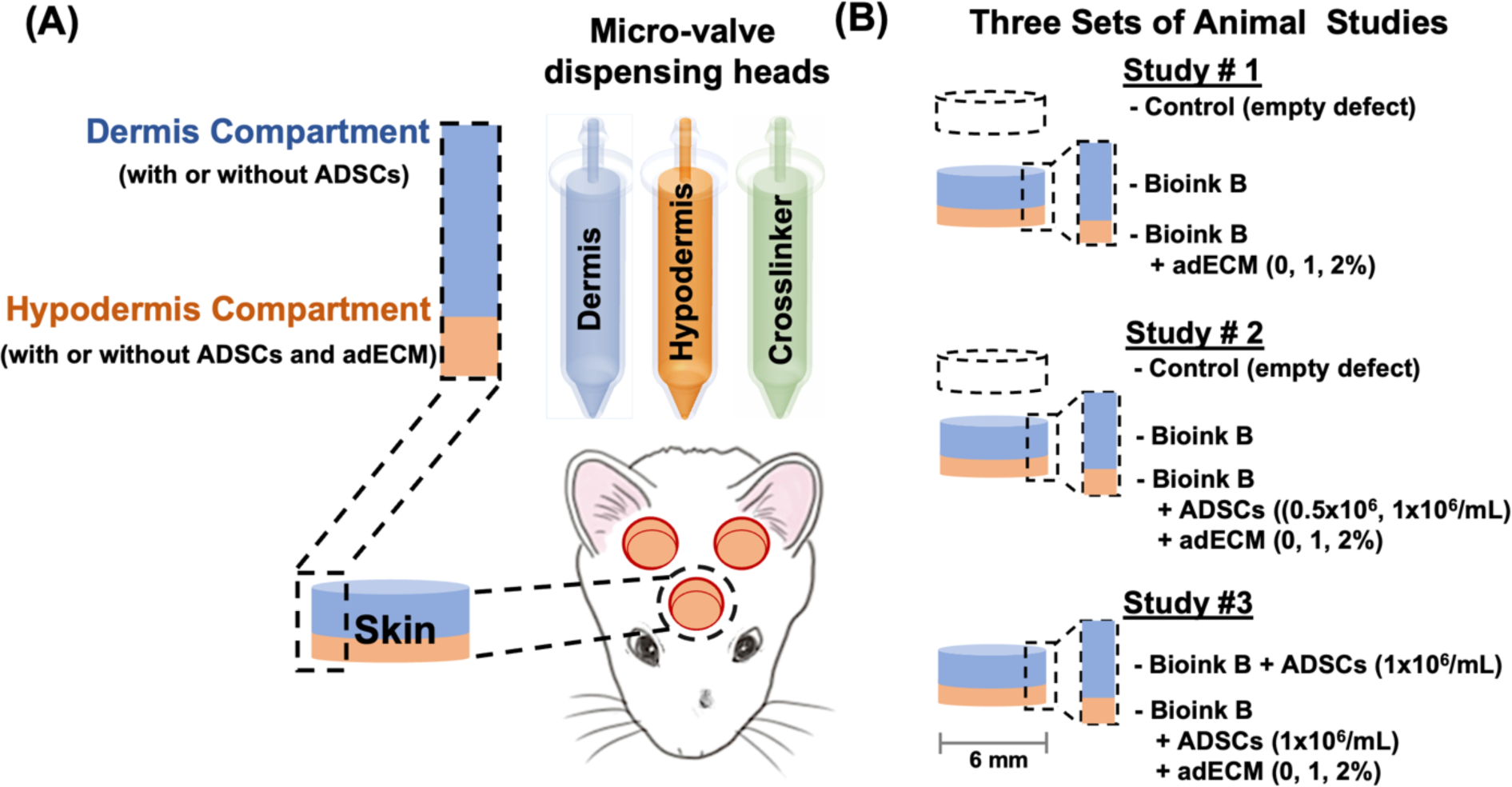
Intraoperative bioprinting (IOB) for full-thickness skin reconstruction. (A) General schematic of the IOB process using droplet-based bioprinting to reconstruct hypodermis and dermis compartments in a surgical setting. IOB was performed on nude rats with three 6-mm full-thickness skin defects on crania for three different sets of animal studies. (B) In Study #1, the role of adECM concentration in the hypodermis compartment on skin reconstruction was evaluated. In Study #2, the combinational role of ADSC and adECM concentration in the hypodermis compartment on skin reconstruction was evaluated. In Study #3, the effect of adECM in adipose tissue formation in ADSC-laden hypodermis and dermis compartments was explored.

## 2. RESULTS AND DISCUSSION

### 2.1 Bioprintability of Human adECM-loaded Bioink

In this study, microvalve-based bioprinting, which is a type of droplet-based bioprinting (DBB) technique, was utilized owing to its advantages of precise control over the deposition rate, droplet size, and the amount of bioink dispersed [33]. For the bioink, human adECM was loaded into fibrinogen at 1 or 2%, which was further supplemented with Factor xiii (Fxiii), also known as fibrin stabilizing factor [34,35]. The rationale for the use of fibrinogen is that it can easily be bioprinted and crosslink with the deposited crosslinker (thrombin) quickly, which enable the gelation of deposited solutions and facilitate a layered structure. In this work, up to 2% adECM was loaded into fibrinogen to demonstate the effectiveness of adECM in inducing adipogenesis in the hypodermis layer even at a very low amount and concentrations over 2% caused cell aggregation, and hence nozzle clogging, during our preliminary work. In our preliminary studies, we also observed that the addition of adECM impaired the crosslinking between fibrinogen and thrombin as adECM was composed of various molecules, such as collagen type I-VII, laminin, fibronectin, elastin, and glycosaminoglycan (GAG), which might dilute the fibrin concentration or interfere with the crosslinking process [36]. Thus, we used Fxiii to strengthen the fibrin clot by catalyzing an active form of thrombin to bond with fibrinogen. As a crosslinker, thrombin was used to polymerize fibrinogen into fibrin, and Fxiii was activated by thrombin to stabilize fibrin[37]. To test the bioprintability of different bioink solutions, including fibrinogen with/without Fxiii and adECM, rheological analysis was performed, where the bioink solutions exhibited similar viscosity profiles without Fxiii but after the addition of Fxiii, the viscosity increased (**Figure 2A**). Next, the jetting behavior using micro-valves was investigated and all bioink solutions revealed similar behavior, where a bulb developed as the tail breaking up into individual droplets (**Figure 2B**). However, all the solutions still showed low viscosity regardless of Fxiii, so the jetting behavior did not differ considerable from each other, which is indicating that the addition of adECM into the bioink up to 2% did not impair its bioprintability.

**Figure 2.**
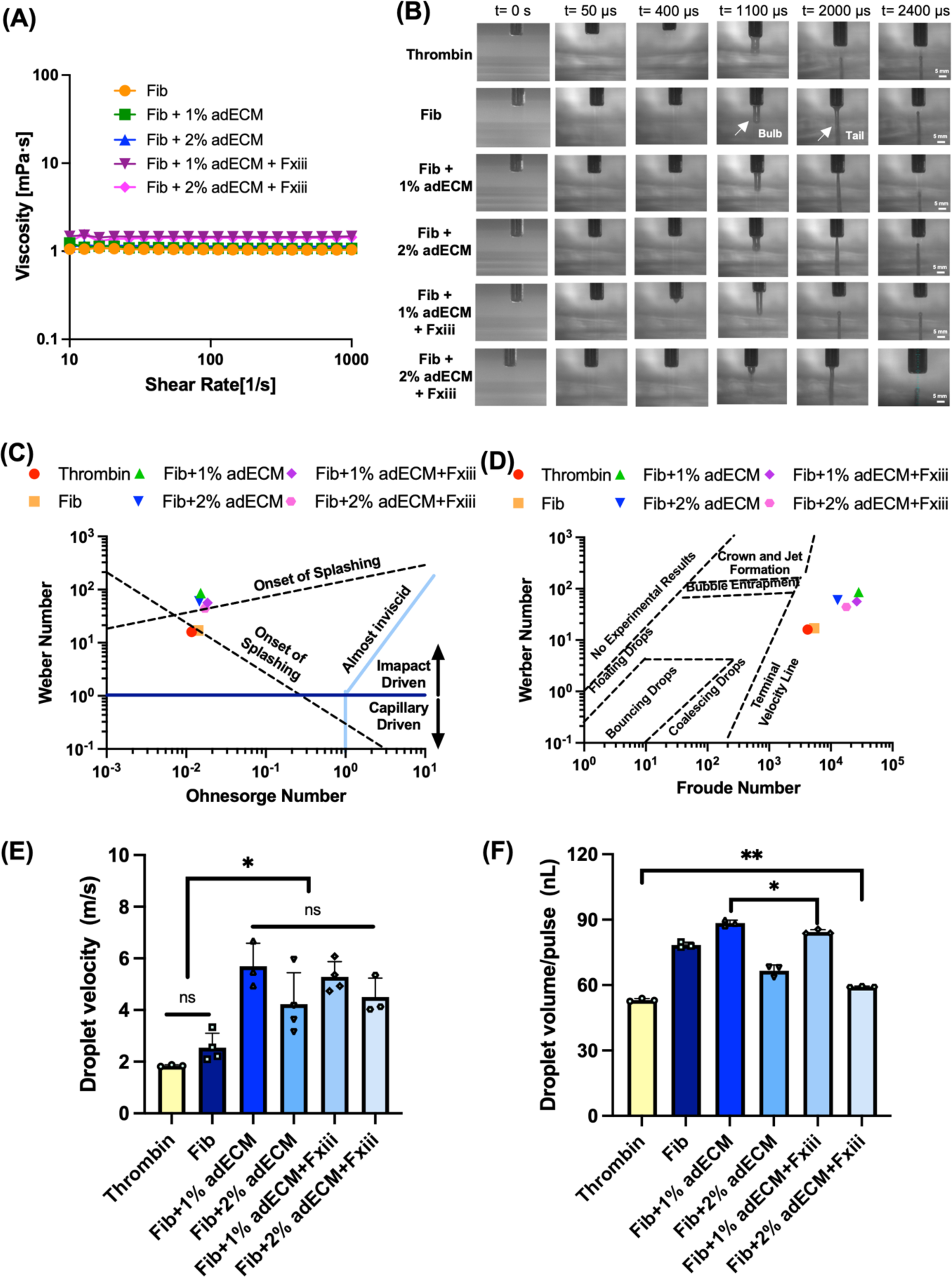
In vitro characterization of bioink solutions. (A) Viscosity of surfactant involved fibrinogen loaded with 0, 1 or 2% (v/v) adECM with and without Fxiii at 25 ℃. (B) Optimization of jetting of bioink solutions, where solutions ejected from the micro-valve device break up into streams of multiple droplets upon exiting the nozzle orifice. (C) Interaction of droplets with a solid substrate, represented by Weber number of droplets as a function of their Ohnesorge number, drawn following the schematic diagram of Schiaffino and Sonin [38]. (D) The interactions of droplets with the liquid substrate, represented by Weber number of droplets as a function of their Froude number, drawn following the schematic diagram of Leng [42]. (E) The droplet velocity (*n*=3) and (F) the droplet volume per pulse (*n*=5) of fibrin solutions with and without Fxiii (p* < 0.05 and p** < 0.05; error bars indicate s.e.m.; ns indicates ‘non significant’; ‘Fib’ indicates fibrinogen).

After droplet formation, the behavior of droplets during their deposition on a solid substrate was evaluated using Reynolds, Weber, Ohnesorge and Bond numbers [38–41]. To be specific, the impact of gravitational forces was negligible since the Bond number of droplets was less than 1. Therefore, either inertial or capillary forces induced the spreading of droplets when they were deposited on the solid substrate. However, spreading of droplets was not significant and bioprinted droplets eventually coalesced into a single droplet due to the resistance of inertial oscillations over the initial impact pressure (**Figure 2C**). After bioprinting of the first layer, the solid-to-liquid interface changed to liquid-to-liquid interface for bioprinted droplets because complete polymerization of fibrin required 10 to 15 min. Here, to describe the liquid-liquid interactions, Froude and Ohnesorge numbers were also utilized [42] (**Figure 2D**). All the solutions were ejected at the terminal velocity, and the addition of adECM significantly increased the droplet velocity (**Figure 2E**) suggesting that the viscous forces (Ohnesorge number) increased, owing to the addition of adECM, but at the same time, the inertia force (Froude number) also increased. Hence, fibrinogen with adECM was ejected at a higher velocity, but the droplet volume decreased due to higher viscosity when adECM concentration increased from 1 to 2% (**Figure 2F**). Nevertheless, those different rheological properties did not cause significant splashing of droplets, so solutions with different compositions were all feasible for bioprinting.

### 2.2 Evaluation of in vitro Bioprinted Constructs

To evaluate the role of Fxiii and adECM on the structural properties of bioprinted constructs, scanning electron microscopy (SEM) images were taken. As shown in **Figure S1A**, Fxiii affected the strength of the fibrin clot by catalyzing an active form of thrombin to bond with fibrinogen. SEM imaging confirmed that fibrinogen crosslinked with thrombin in the absence of Fxiii consists of a loose, turbid and coarse network while fibrinogen crosslinked with thrombin in the presence of Fxiii consists of a thick and dense network of significantly thicker fibrils. The addition of adECM either with or without Fxiii showed a dense and compact network with thinner fibrils. We then tested the cytocompatibility of constructs by loading ADSCs at a concentration of 1 million cells per mL. Cell viability after bioprinting, both in the absence and presence of Fxiii, was greater than 95% (**Figures S1B**). Furthermore, the viability of ADSCs was also found to be greater than 95% at Days 1 and 7 (**Figure S2**). Overall, the use of adECM and the bioprinting process did not impair cell viability.

We then tested the adipogenic potential of adECM-loaded bioink, where 3D bioprinted ADSC-laden constructs were maintained in adipogenic differentiation or growth medium. The constructs containing adECM enabled ADSCs to differentiate into adipocytes at Days 14 and 28 in differentiation or growth medium while fibrin-only constructs stained negative to Oil Red O (**Figure 3A**), indicating that adECM had de novo capabilities in supporting adipogenic differentiation. Furthermore, 2% adECM-containing constructs showed stronger Oil Red staining compared to that of 1%. In adECM-containing constructs, stronger Oil Red staining was observed at Day 28 compared to Day 14. We did not see a significant difference among samples cultured with the differentiation media versus growth media on both Days 14 and 28.

**Figure 3.**
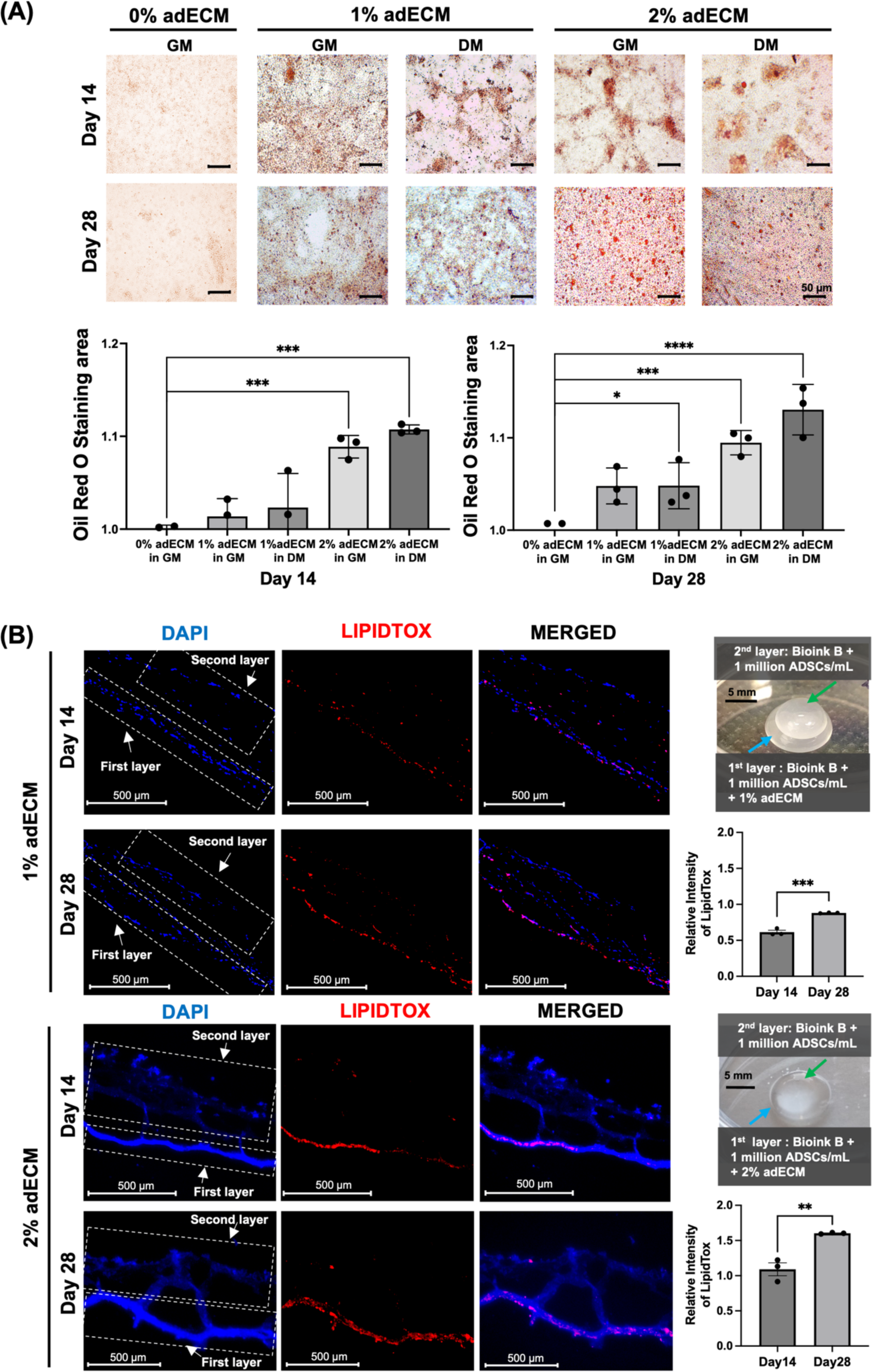
In vitro adipocyte differentiation of ADSCs in 3D bioprinted constructs: (A) Oil red staining for evaluation of adipogenic differentiation of ADCSc (1 million/mL) in the presence of differentiation medium (DM) or growth medium (GM) at Days 14 and 28 (*n*=3; p* < 0.05, p*** < 0.001, p****< 0.0001). (B) LipidTox staining of dual-layer bioprinted constructs, where the hypodermis compartment was loaded with Bioink B, 1 or 2% adECM, and 1 million/mL ADSCs, and the dermis compartment was loaded with Bioink B and 1 million/mL ADSCs in the absence of differentiation media at Day 14 and 28 (*n*=3; p** < 0.01, p*** < 0.001).

We then bioprinted multi-layered constructs with two compartments, where the based bioink solution (henceforth named “Bioink B”) optimized in the presented in vitro study was utilized. The hypodermis compartment (Bioink B loaded with ADSCs at a density of 1 million cells per mL and adECM at a concentration of 1 or 2%) was bioprinted first followed by the dermis compartment (Bioink B loaded with ADSCs at a density of 1 million cells per mL) on top. Constructs were cultured in growth medium for 14 or 28 days and then we evaluated the formation of adipocytes via LipidTox staining (**Figure 3B**). We observed that the hypodermis compartment showed substantially higher numbers of lipid droplets compared to the dermis compartment. Moreover, 2% adECM supported superior differentiation of ADSCs into adipocytes compared to 1% adECM. For both 1 and 2 % adECM, the LipidTox intensity increased from Day 14 to 28.

These findings suggest that the adECM-containing bioink exhibited adipogenic properties and supported the differentiation of ADSCs into adipocytes when cultured in the growth medium, which was further escalated when treated with the differentiation medium.

### 2.3 Intraoperative Bioprinting for CMF Full-thickness Skin Reconstruction

Herein, we demonstrated IOB for the reconstruction of full-thickness skin defects in the CMF zone of 10-12 weeks old female immune-deficient rats. To demonstrate adipogenic differentiation of ADSCs into adipocytes and formation of an adipose layer, we used female rats as they possess a substantial fat component in their skin compared to their male counterparts [43]. In accordance with the principles of the 3Rs (Replacement, Reduction and Refinement) [44], we opted to have three defects per animal thereby reducing the overall number of rats used, concurrently ensuring sufficient distance between defects for clear analysis. Three 6-mm-wide circular skin defects were created on the calvaria of each rat. As shown in **Figure 1B**, three different sets of animal studies were performed. The reader is referred to **Table 1** for details pertaining animal study groups. For IOB, we utilized the micro-valve bioprinting system to eject bioink droplets with high controllability of droplet size and reproducibility for the compartments of skin layers, as reported previously [45]. Using the base bioink solution (Bioink B), we prepared two bioinks, one for the hypodermis (adipose) layer and the other for the dermis layer. The process of IOB took from 5 to 20 min per rat depending on the group type. After the surgery, the wound sites were photographed every day until Day 10 and after that every other day until Day 28.

**Table 1.** In vivo study groups for IOB of the skin tissue.

| Studies |  | Hypodermis |  | Dermis | End Point (weeks) | Total # of defects |
| --- | --- | --- | --- | --- | --- | --- |
|  |  | ADSCs (million/mL) | adECM (%) | ADSCs (million/mL) |  |  |
| Negative control |  | 0 | 0 | 0 |  | 6 |
|  |  | 0 | 0 | 0 |  | 6 |
|  |  | 0 | 0 | 0 |  | 6 |
| Study #1 | G1 | 0 | 0 | 0 |  | 6 |
|  | G2 | 0 | 1 | 0 |  | 6 |
|  | G3 | 0 | 2 | 0 |  | 6 |
| Study #2 | G4 | 0.5 | 0 | 0 |  | 6 |
|  | G5 | 0.5 | 1 | 0 |  | 6 |
|  | G6 | 0.5 | 2 | 0 |  | 6 |
|  | G7 | 1 | 0 | 0 |  | 6 |
|  | G8 | 1 | 1 | 0 |  | 6 |
|  | G9 | 1 | 2 | 0 |  | 6 |
| Study #3 | G10 | 1 | 0 | 1 |  | 6 |
|  | G11 | 1 | 1 | 1 |  | 6 |
|  | G12 | 1 | 2 | 1 |  | 6 |

In Study #1, the hypodermis compartment was bioprinted with Bioink B loaded with different concentrations of adECM including 0% (G1), 1% (G2), and 2% (G3) in order to evaluate the role of adECM in formation of adipose tissue in the hypodermis layer (**Figure 4A and Video S1**). After bioprinting of the hypodermis compartment, the dermis was bioprinted using Bioink B. As the defects were not substantially large, the epidermis compartment was not bioprinted as keratinocytes were expected to migrate over the bioprinted constructs. Greater wound closure of ∼80, ∼82, and ∼84% was observed in G1, G2, and G3, respectively, as compared to the control group (∼72%) at Day 14. In addition, G1, G2 and G3 represented faster re-epithelialization of ∼72, ∼73, and ∼70% at Day 7, respectively, compared to the control group (∼40%) (**Figures 4B and S3**). Furthermore, Hematoxylin and Eosin (H&E) staining demonstrated the overall structure of repaired wounds at Day 14 (**Figure 5A**) and Day 28 (**Figure S4B**). At Week 2, bioprinted groups showed organized connective tissue with a well-defined dermis layer, while the control group showed less re-organized and remodeled structure. Interestingly, we found hair follicle-like downgrowths in the dermis particularly adECM containing groups (**Figure 5B**). Compared to the control group, G2 and G3 showed ∼3 downgrowths on average, while G1 showed 1, as observed from H&E images in **Figures 5A**. This indicates that adECM supported the growth of hair follicle-like structures, which could be due to the fact that the addition of the adECM niche promoted adipogenesis, leading to section of BMP-2, which helped activate hair follicle stem cells [24]. Furthermore, Masson’s Trichrome staining (MTS) was conducted to demonstrate collagen deposition and vascularization at Week 2 (**Figures 5A and C**) and Week 4 (**Figure S4B**). According to the collagen index determined from the MTS images, bioprinted groups represented more densely packed collagen fibers as opposed to the control group. To explore the role of adECM in formation of adipocytes and deposition of lipid droplets in hypodermis, we performed LipidTox staining. G2 and G3 exhibited stronger LipidTox staining but not statistically significant, compared to the control and G1, demonstrating that bioprinting adECM-containing bioink facilitated adipogenesis at Days 14 and 28 (**Figures 5A, D and S4C**). Blood vessels in all groups were also identified as highlighted with orange arrows in MTS images (**Figure 5A**). Based on CD31 staining images (**Figure S4A**), we observed vascularization in all groups confirming the findings from MTS, where blood vessels were detected in all groups. Particularly, CD 31 expression showed higher in G3 compared to G1 as shown in **Figure 5E**. Overall, the results suggests that the addition of adECM into the bioink enhanced adipogenesis and vascularization in hypodermis and facilitated not only full thickness skin reconstruction but also the downgrowth of hair follicle-like structures.

**Figure 4.**
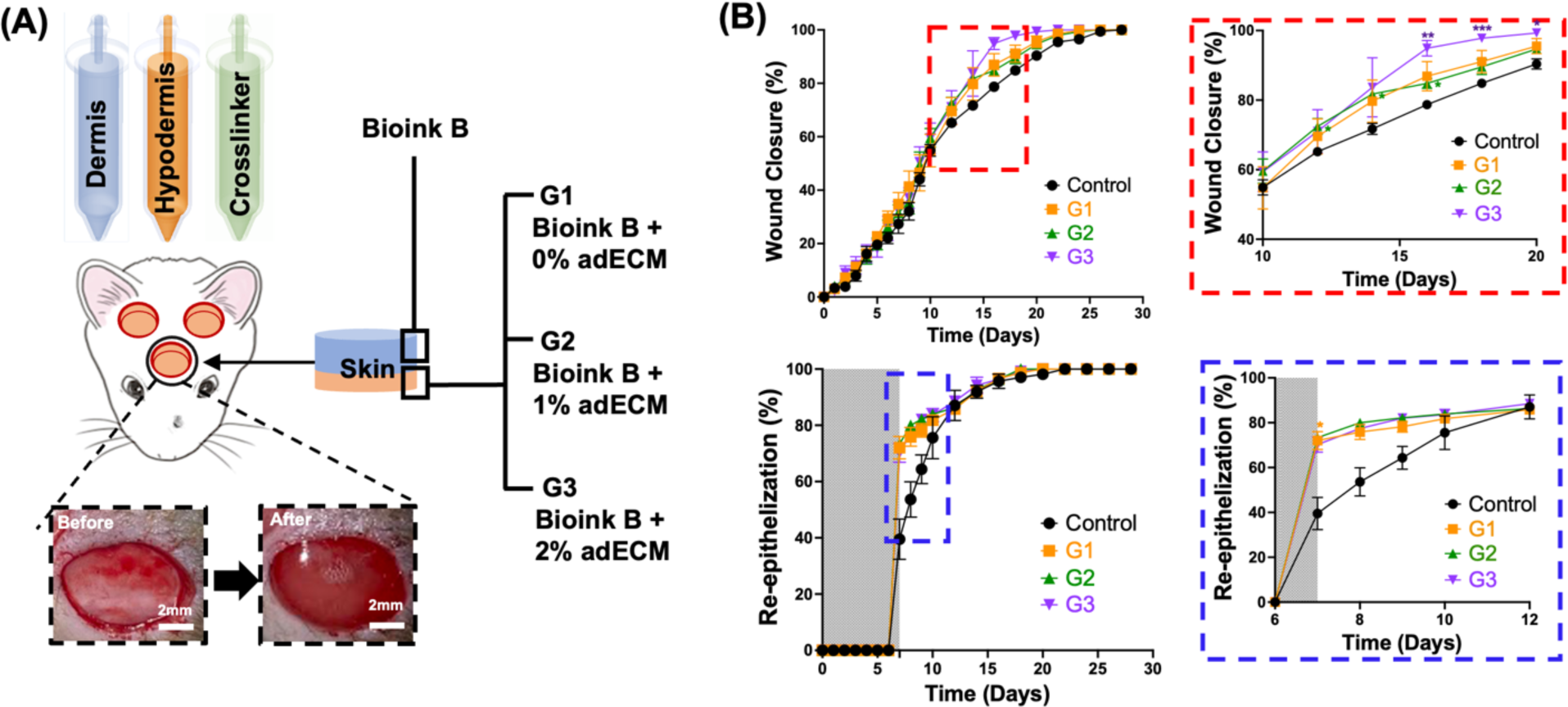
Study #1: (A) A schematic of IOB experiments for decoding the role of presence of adECM in the hypodermis layer. IOB was performed on nude rats with three 6-mm full-thickness skin defects on crania for a total of four groups including i) the negative control group (empty defects), ii) Bioink B + 0% adECM (G1), iii) Bioink B + 1% adECM (G2), and iv) Bioink B + 2% adECM (G3), where Bioink B was bioprinted on top of the hypodermis compartment. (B) Percentage (%) wound closure (top) and re-epithelization (bottom) (*n*=6; p*<0.05, p**<0.01, p***<0.001; color code shows the difference with respect to the control).

**Figure 5.**
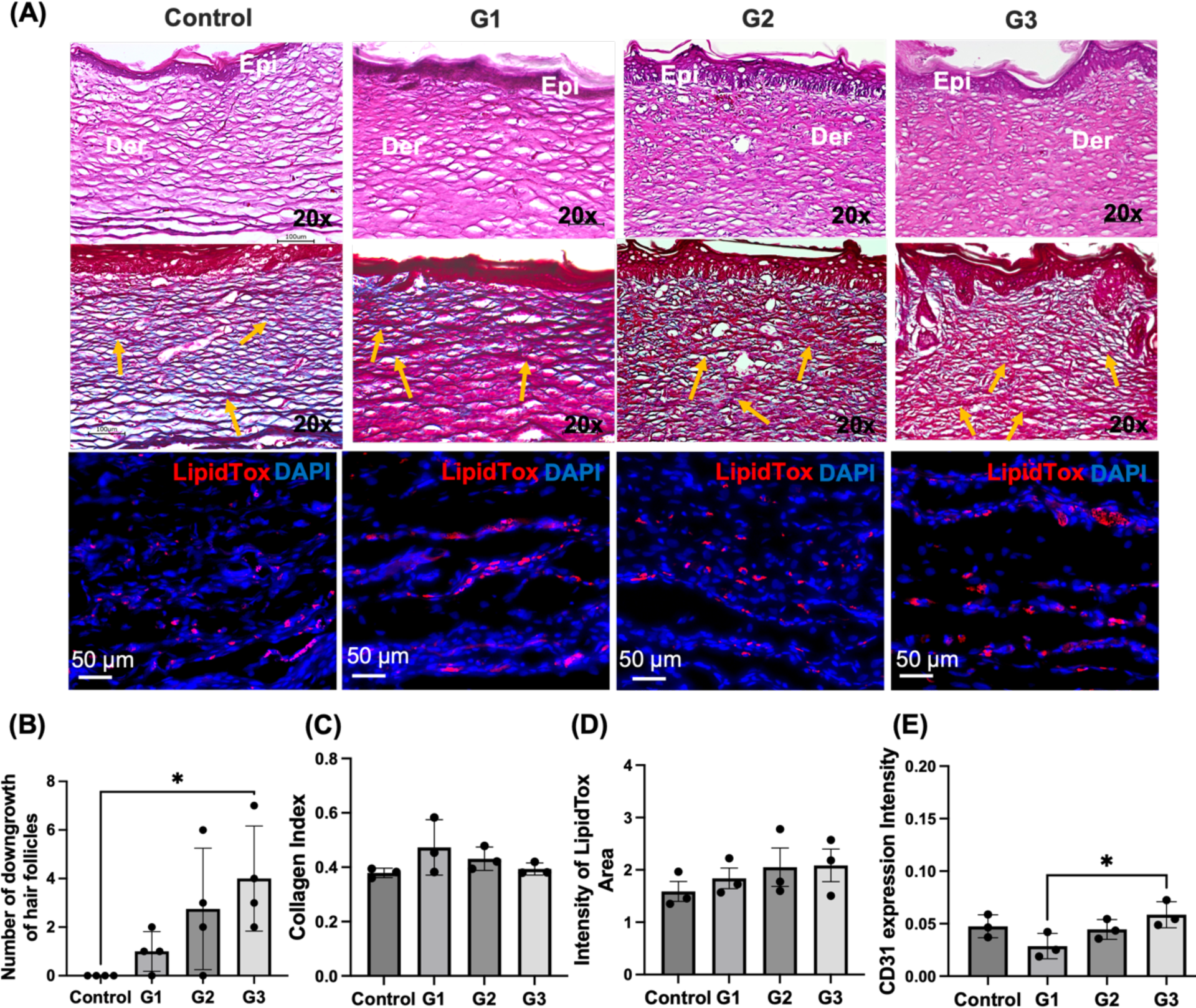
(A) Histological images of the reconstructed skin stained for H&E (Epi: epidermis layer; Der: dermis layer), MTS (orange arrows indicate blood vessels), LipidTox at Day 14. (B) The number of hair follicle downgrowth quantified using H&E images (*n*=3; p*< 0.05). (C) Collagen index determined from MTS images (*n*=3). (D) Fluorescence intensity of LipidTox (*n*=3). (E) Expression of CD31 from immunostaining images (*n*=3; p* < 0.05).

We then performed Study #2 to understand the combinational role of adECM and ADSCs in the reconstruction of full-thickness skin defects. IOB using Bioink B with different concentrations of adECM and densities of ADSCs, including 0.5 million per mL ADSC-laden Bioink B with 0% adECM (G4), 1% adECM (G5) and 2% adECM (G6), and 1 million per mL ADSC-laden Bioink B with 0% adECM (G7), 1% adECM (G8) and 2% adECM (G9) (**Figure 6A, Videos S2 and S4**). Empty defects were used as a control group. After bioprinting of the hypodermis compartment, the dermis compartment was bioprinted with Bioink B. Morphometric evaluation of wound healing was performed by calculating the percentage of wound closure and wound re-epithelialization with respect to the original wound using the same approach described in Study #1 (**Figure 6B**). Based on the wound photographs presented in **Figure S5**, wound closure (%) and re-epithelialization (%) was evaluated. Compared to the control group at Day 14 (∼75%), all ADSC-laden Bioink B groups showed improved wound closure. Particularly, the groups containing 1 million ADSCs per mL, namely G7, G8, and G9, showed faster wound closure of ∼90, ∼91, and ∼88%, respectively, compared to 0.5 million per mL ADSC-laden Bioink B groups, where G4, G5, and G6 showed wound closure of ∼89, ∼85, and ∼85%, respectively. Similar to wound closure results, all ADSC-laden groups (G4, G6, G6, G7, G8 and G9) exhibited faster re-epithelialization, namely ∼69, ∼74, ∼74, ∼61, ∼70, and ∼71% at Day 7, compared to the control group ∼45%, respectively. The wound closure was accelerated when ADSCs were bioprinted, where ADSCs are known to have the potential to differentiate into various lineages such as fibroblasts, keratinocytes, and endothelial cells to enhance wound healing and angiogenesis, release growth factors and cytokines, and possess antibacterial properties to accelerate the wound healing process [46].

**Figure 6.**
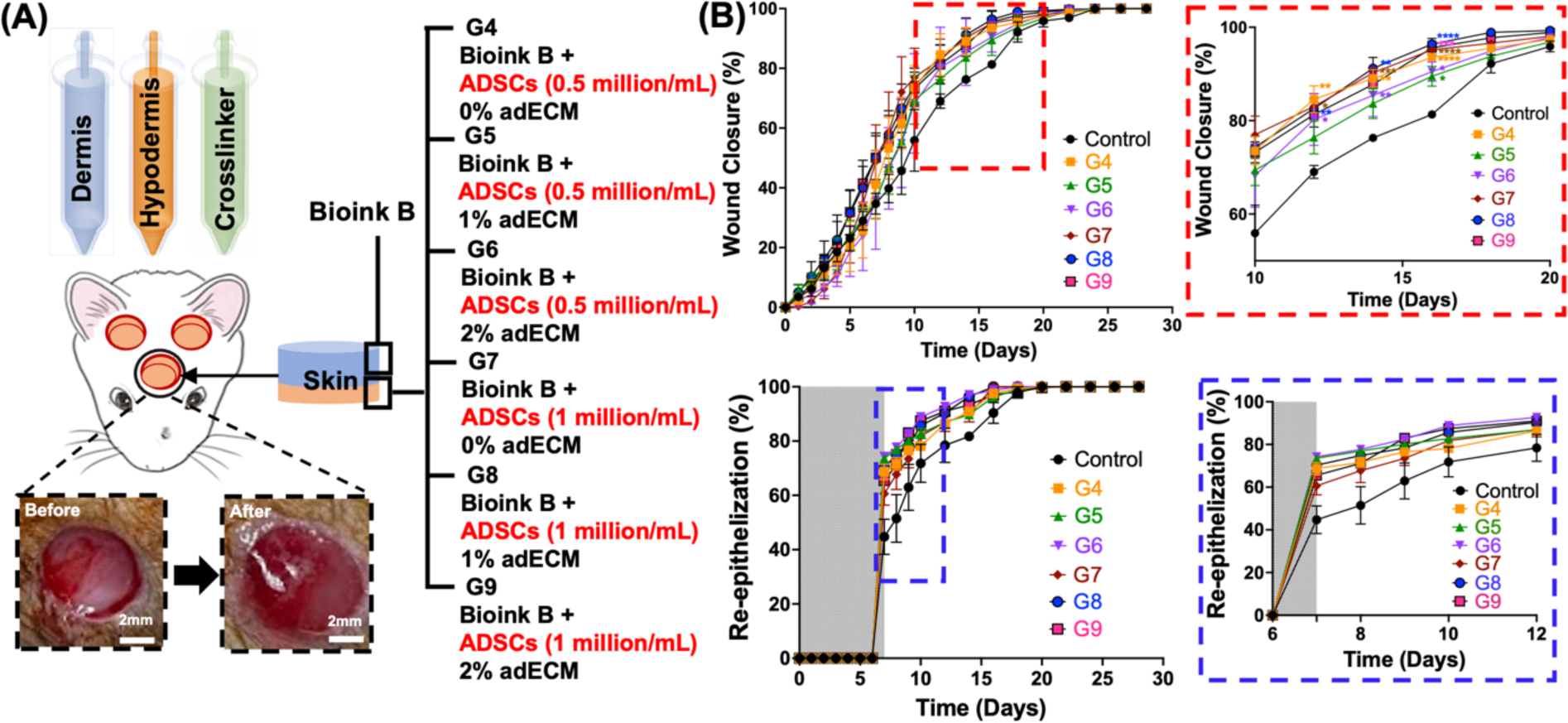
Study #2: (A) A schematic of IOB experiments for decoding the combinational role of adECM and ADSCs in reconstructing the adipose tissue in the hypodermis compartment. IOB was performed on nude rats with three 6-mm full-thickness skin defects on crania for a total of seven groups including i) the negative control group (empty defects), ii) Bioink B + 0.5 million/mL ADSCs + 0% adECM (G4), iii) Bioink B + 0.5 million/mL ADSCs + 1% adECM (G5), iv) Bioink B + 0.5 million/mL ADSCs + 2% adECM (G6), v) Bioink B + 1 million/mL ADSCs + 0% adECM (G7), vi) Bioink B + 1 million/mL ADSCs + 1% adECM (G8), and vii) Bioink B + 1 million/mL ADSCs + 2% adECM (G9), where Bioink B was bioprinted on top of the hypodermis compartment. (B) Percentage (%) wound closure (top) and re-epithelization (bottom) (*n*=6; p*<0.05, p**<0.01, p***<0.001, p****<0.0001; color code shows the difference with respect to the control).

To further evaluate the reconstructed skin over time, we conducted histological analysis at Week 2 (**Figure 7**) and Week 4 (**Figure S6**). H&E images demonstrated that the epidermis and dermis layers in all bioprinted groups were organized and reconstituted in two weeks compared to the control group. Similarly, MTS showed that collagen deposition in all bioprinted groups was slightly more apparent compared to the control group at Week 2. More importantly, there was a significantly distinctive hair follicle downgrowths in 1 million per mL ADSC-laden Bioink B groups (G7, G8, and G9) compared to 0.5 million per mL ADSC-laden Bioink B (G4, G5, and G6) and the control groups. As highlighted in **Figure 7B**, the average number of hair follicle downgrowth for G7, G8 and G9 were 3, 5, and 5, respectively, which was higher than the other groups. This is suggestive that the bioprinted ADSCs and adECM niche might enhance adipogenesis, which has a critical role in hair follicle cycling, particularly extending the anagen stage of hair growth [47]. From the collagen index perspective, G4, G5, G6, G7, G8 and G9 showed slightly more collagen deposition compared to control group, but not significantly different except G8 (**Figure 7C**). To further support our findings, we performed LipidTox assay which demonstrated that the co-delivery of ADSCs and adECM (G5, G6, G8 and G9) exhibited stronger expression of LipidTox staining and hence better differentiation of adipocytes as compared to G2 and G3 in the first set of animal study, implying that adECM and ADSCs together supported adipogenesis during the process of wound healing (**Figures 7D and S6C**). Based on CD31 staining images (**Figures S6A**), we observed vascularization in all groups confirming the findings from MTS (**Figure 7A**), where blood vessels were detected in all groups. G8 and G9 showed higher intensity of CD31 expression, implying stronger vascularization compared to other groups (**Figure 7E**).

**Figure 7.**
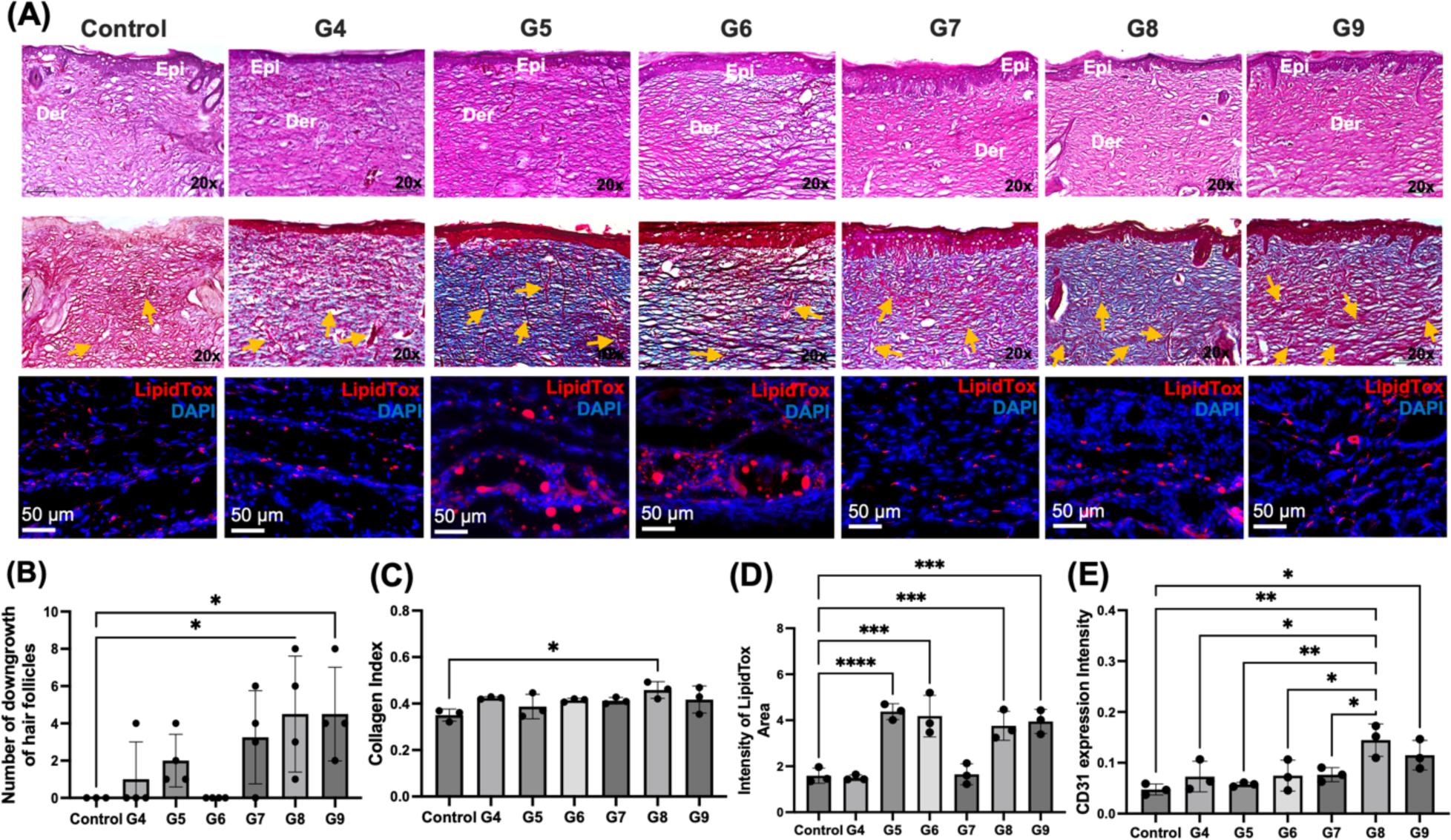
(A) Histological images of the reconstructed skin stained for H&E (Epi: epidermis layer; Der: dermis layer), MTS (orange arrows indicate blood vessels), LipidTox at Day 14. (B) The number of hair follicle downgrowth quantified using H&E images (*n*=3; p* < 0.05). (C) Collagen index determined from MTS images (*n*=3; p* < 0.05). (D) Fluorescence intensity of LipidTox (*n*=3; p*** < 0.001, p****< 0.0001). (E) Expression of CD31 from immunostaining images (*n*=3; p* < 0.05, p** < 0.01).

In Study #3, we explored the role of adECM in skin reconstruction when ADSCs were bioprinted in both hypodermis and dermis compartments (**Figure 8A and Video S4**). The density of ADSCs yielding the highest level of adipogenesis, vascularization, and the fastest wound healing from the previous sets of animal studies were utilized. Similar to Study #1 and #2, using the wound photographs (**Figure S7**), skin repair was morphometrically evaluated by measuring the percentage of wound closure and re-epithelialization with respect to the original wound (**Figure 8B**). Overall, the wound closure for G10 (∼91%), G11 (∼93%), and G12 (∼91%) at Day 14 were slightly higher than those for Study #1 and #2 (discussed previously) while the re-epithelization rates for G10 (∼66%), G11 (∼62%), and G12 (∼65%) at Day 7 were slightly lower than those for Study #1 and #2. To evaluate the histomorphology of reconstructed skin over time, we performed histological assays at Week 2 (**Figure 9A**) and Week 4 (**Figure S8**). H&E images showed well-formed skin structures in all groups. More interestingly, G10, G11, and G12 exhibited higher number of hair follicle downgrowth on average compared to the groups presented in Study #1 and #2 (**Figures 9B**). Among studies #1, #2 and #3 with 2% adECM (G3, G6, G9, and G12), the G12 group showed obvious orientation of downgrowths (**Figure S9**). M&T images demonstrated that G10, G11, and G12 had similar collagen deposition (**Figure 9C**). LipidTox staining demonstrated that G11 and G12 showed higher intensity of LipidTox compared to G10 and other groups in the previous studies at Day 14 and 28 as well (**Figures 9D and S8C**). Additionally, we observed a similar level of vascularization in all groups, which also confirmed our findings with MTS and CD31 immunostaining images (**Figures 9E and S8A**). We further confirmed adipocyte differentiation by Zsgreen transfected ADSCs from GFP staining and FABP4 (Fatty acid binding protein 4) staining in the hypodermal compartment at Day 14. Though we bioprinted tdTomato transfected ADSCs in the dermis compartment after bioprinting Zsgreen transfected ADSCs in the hypodermis, we hardly observed tdTomato^+^ ADSCs. We speculate that the replacement of Tegaderm^TM^ transparent dressings to cover the bioprinted sites every one or two days to maintain wound moisture, hygiene and drainage in the wound might cause the loss of some of the bioprinted construct in the dermal compartment. Compared to G10 and G11, we obviously observed adipocyte differentiation from GFP and FABP4 staining in G12, indicating that bioprinted Zsgreen^+^ ADSCs were differentiated into adipocytes in the hypodermis compartment (**Figure 9F**). Although further biological assessements, such as single cell RNA sequencing and western blot, will support our findings, these were beyond the scope of this study and we aim to explore molecular-scale features of evolution of skin structure over time in a future work. Therefore, taken all together into account, the last group, G12, seems that 2% adECM with ADSCs at a density of 1 million cells per mL greatly enhance adipogenesis and facilitating the downgrowth of hair follicle-like structures.

**Figure 8.**
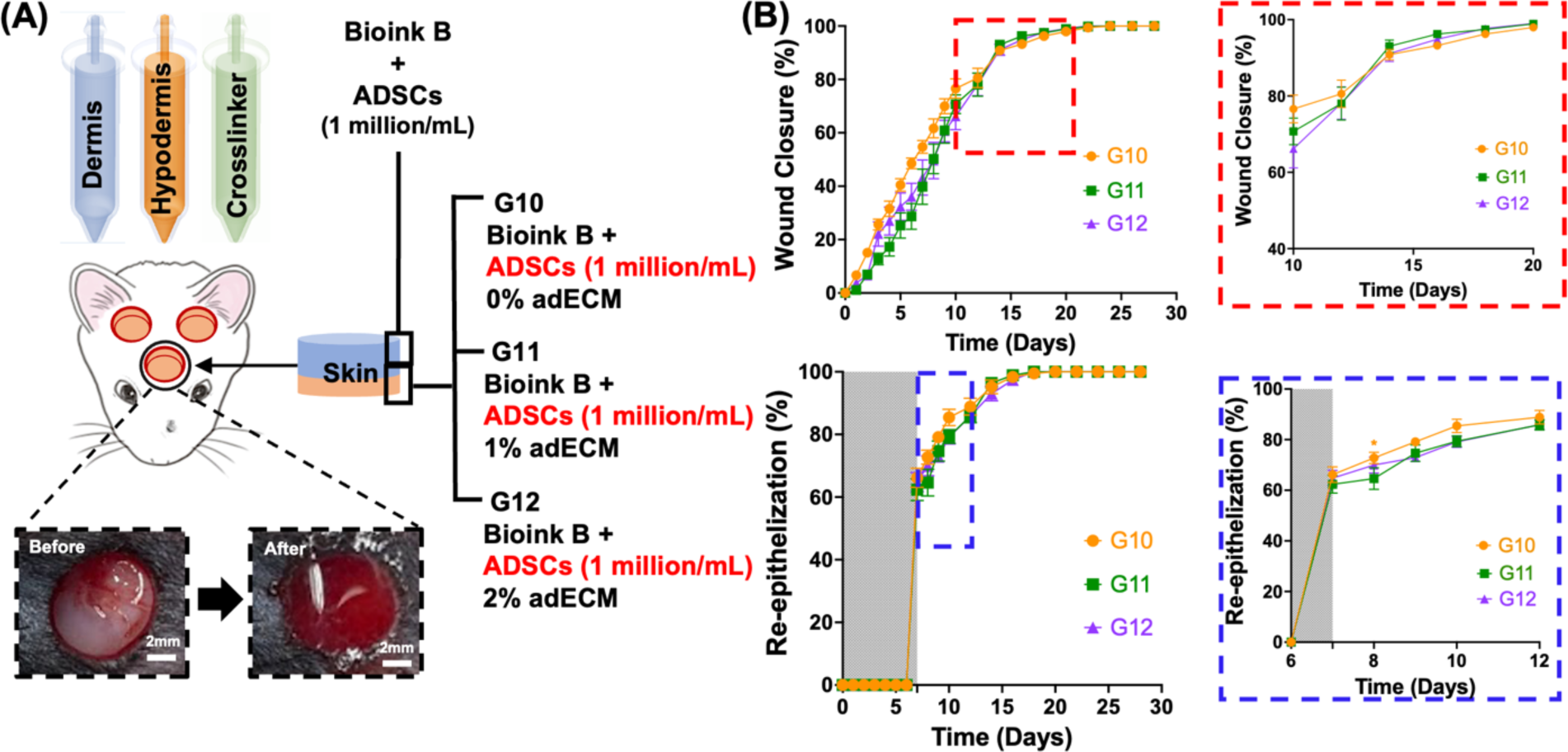
Study #3: (A) A schematic of IOB experiments for decoding the role of adECM in adipose tissue formation when ADSCs were laden in hypodermis and dermis compartments. IOB was performed on nude rats with three 6-mm full-thickness skin defects on crania for a total of three groups including i) Bioink B + 0% adECM (G10), ii) Bioink B + 1% adECM (G11), iii) Bioink B + 2% adECM (G12), where Bioink B + 1 million/mL ADSCs were bioprinted in the dermal compartments for all groups. (B) Percentage (%) wound closure (top) and re-epithelization (bottom) (*n*=6; p*<0.05; color code shows the difference with respect to G12).

**Figure 9.**
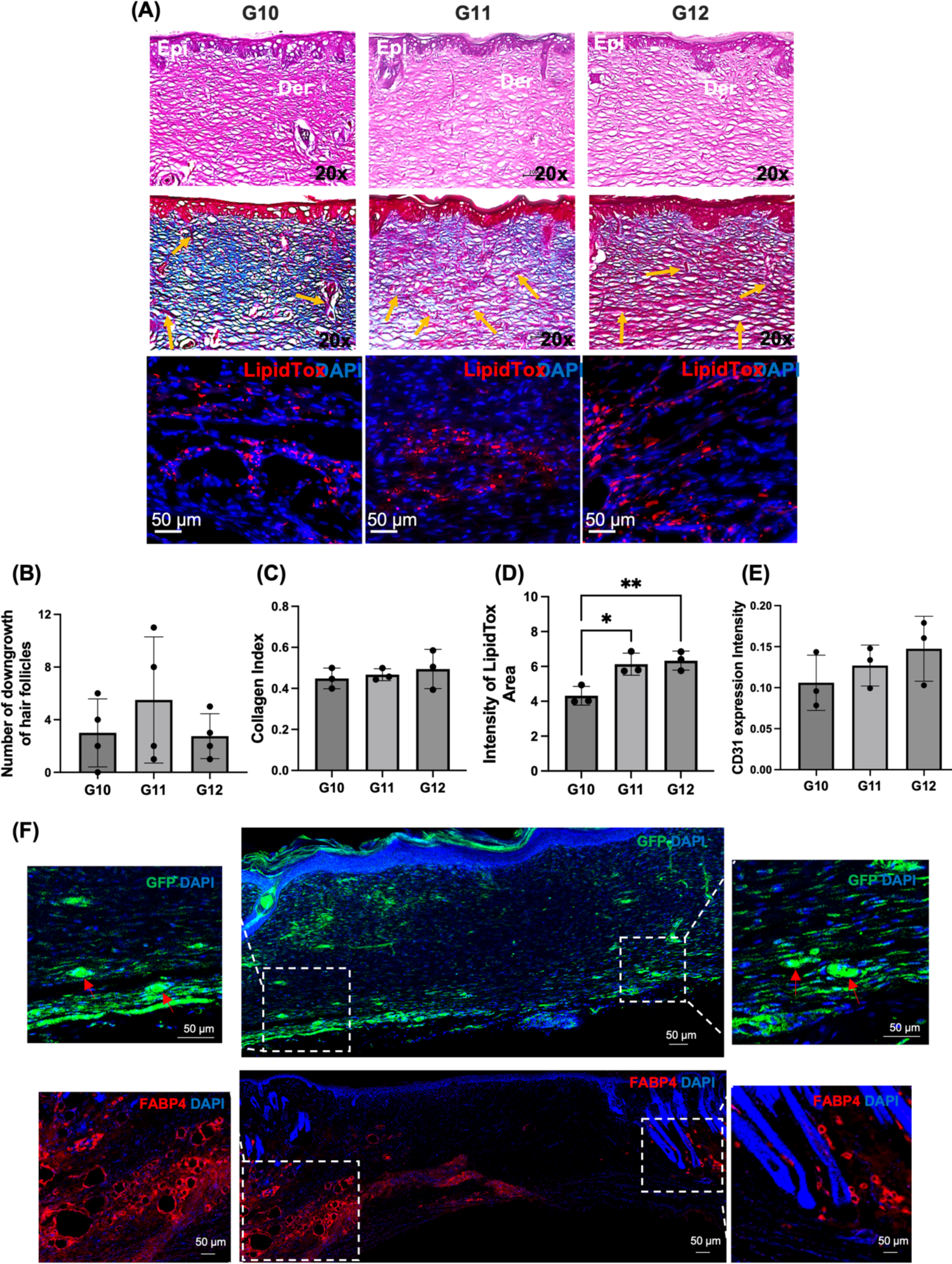
(A) Histological images of the reconstructed skin stained for H&E (Epi: epidermis layer; Der: dermis layer), MTS (orange arrows indicate blood vessels), LipidTox at Day 14. (B) The number of hair follicle downgrowth quantified using H&E images (*n*=3). (C) Collagen index determined from MTS images (*n*=3). (D) Fluorescence intensity of LipidTox (*n*=3; p* < 0.05, p** < 0.01). (E) Expression of CD31 from immunostaining images (*n*=3). (F) Histological images of the reconstructed skin stained for DAPI and GFP-fluorescence distribution with ADSCs transfected with Zsgreen lentivirus and for DAPI and FABP4 at Day 14.

## 3. CONCLUSION

In this work, we demonstrated the reconstruction of CMF full-thickness skin defects using the IOB technology, where a new bioink was formulated using adECM obtained from clinical lipectomy samples. The presented approach accelerated the reconstruction of full-thickness skin defects and facilitated up to near-complete wound closure in two weeks. More importantly, the combinatorial use of ADSCs and adECM enabled the formation of adipocytes in the hypodermal compartment. Interestingly, adECM in conjunction with adipogenesis increased the formation of downgrowth’s of structures that resembled hair follicles. Although adipocytes do not directly contribute to the cellular structure of hair follicles, they are important for homeostasis and maintenance [48]. In our experiments, adipocytes may have altered the ECM to be more supportive for downgrowth formation. With the fully automated bioprinting ability and compatible materials at the clinical grade, the IOB technology will have a significant impact on the clinical translation of precisely reconstructed skin in the near future.

## ACKNOWLEDGEMENT

This work has been supported by National Institute of Dental and Craniofacial Research (NIDCR) Award R01DE028614 (I.T.O.), National Institute of Arthritis and Musculoskeletal and Skin Diseases (NIAMS) Award R21AR082668 (I.T.O.) and R01AR078743 (R.R.D.), and 2236 CoCirculation2 of TUBITAK award 121C359 (I.T.O.). We thank Dr. Daniel Hayes’ laboratory (Penn State) for preparation of adECM, Dr. Srinivas Koduru (Penn State) for processing fat tissue, Dr. Dishary Banerjee (Penn State) for her assistance in animal surgeries and Dr. Sepehr Nesaei (Penn State) for his assistance with pilot micro-valve bioprinting studies. The opinions, interpretations, conclusions, and recommendations are those of the author and are not necessarily endorsed by NIDCR, NIAMS, and TUBITAK.

## DATA AVAILIBILITY

All data needed to evaluate the conclusions in the paper are present in the paper and the supporting information. Additional data related to this paper may be requested from the authors.

## 4. MATERIALS AND METHODS

### 4.1 Cell Culture

ADSCs were purchased from Lonza (Walkersville, MD) and cultured in Dulbecco’s modified Eagle medium (DMEM) supplemented with 20% fetal bovine serum (FBS, Biotech, Minneapolis, MN) and 1% penicillin/streptomycin (Sigma-Aldrich, St. Louis, MO) at 37 °C and 5% CO_2_ in an incubator. The medium was changed every 2 or 3 days.

For the transduction of ADSCs with *Zoanthus sp.* green fluorescent protein (ZsGreen) and Tandem-dimer tomato (tdTomato) lentivirus, ADSCs were transduced at passages 4-5 (∼50% confluency) with EF1 ZsGreen lentiviral vector (Vectalys, Toulouse, France) and EF1 tdTomato lentiviral vector (Vectalys). A multiplicity of infection (MOI) of 20 was maintained for the transduction process. Briefly, a transduction mix was prepared by adding a measured amount of viral vector solution in complete culture media and 800 µg/ml polybrene (Sigma). The transduction mix was then transferred to a flask at ∼50% confluency and incubated for 8 h. The transduction mix was then discarded, and the flask was rinsed with Dulbecco’s Phosphate Buffered Saline (DPBS) (1X) and replenished with culture media. Cells were then allowed to grow in flasks for another 48 h before sorting them on a MoFlo Astrios sorter (Beckman Coulter, Pasadena, CA) for the brightest cells. The brightest cells were then collected for further cell culture.

### 4.2 Adipose Tissue Procurement and Processing to Obtain Human adECM

Decellularized ECM was prepared as described in the previous work [49]. Briefly, adipose tissue samples were collected from selective patients with consent, receiving panniculectomy surgery under the protocol approved by the Institutional Review Board (STUDY 00014177). To ensure confidentiality, the donors’ information was de-identified for all specimens. Excised adipose tissue was mechanically disrupted and enzymatically digested with collagenase and vessels were excised from the adipose tissue for ECM extraction. Fibrous tissue that could not be mechanically homogenized was continuously removed and collected, while all other tissue components were discarded. Urea buffer was added to the collected fibrous tissue and the solution was then centrifuged. The pelleted tissue was collected, dialyzed, and then digested with pepsin. The solution was dialyzed against deionized water for a day before being lyophilized, yielding adECM powder. The adECM power was sterilized with ethylene oxide gas.

### 4.3 Preparation of Fibrin-Based Bioink for Dermal and Hypodermal Compartments

Fibrinogen solution, the base bioink (Bioink B), was prepared at a concentration of 5 mg/mL by dissolving fibrinogen protein (crystals, Sigma-Aldrich) in sterile phosphate-buffered saline (PBS) (VWR) and stored at −30 °C for future use. Similarly, thrombin stock solution was prepared at a concentration of 2 U/mL by dissolving thrombin powder (Sigma-Aldrich) in PBS and stored at −30 °C for future use. Coagulation Factor XIII (Fxiii) was stored at −30 °C and used at 1 g/ L in 10 mM of calcium chloride (CaCl_2_) solution (Sigma-Aldrich). For the crosslinker, 2 U/mL thrombin was mixed with 1 mg/mL Fxiii (in 10 mM CaCl_2_). To fabricate full-thickness skin tissue constructs, Bioink B was loaded with adECM and/or ADSCs. For the hypodermal compartment, adECM containing Bioink B combined with fibrinogen and ZsGreen^+^ ADSCs were prepared and bioprinted along the thrombin solution as a crosslinker. The final concentration of the bioink was 1 or 2% (w/v) adECM and 5 mg/mL fibrinogen. Along with that, the concentration of ZsGreen^+^ ADSCs was 0.5 or 1 million/mL. For the dermal compartment, Bioink B was loaded with tdTomato^+^ ADSCs without adECM, where the concentration of tdTomato^+^ ADSCs was 1 million/mL.

### 4.4 Rheology Study

To investigate the rheological behavior of solutions, including fibrinogen, fibrinogen with 1 or 2% adECM and fibrinogen loaded with Fxiii and 1 or 2% adECM, flow sweep measurements were conducted with a rheometer (MCR 302, Anton Paar, Austria) in the shear rate range of 10– 1000 s^−1^. Measurements were performed at room temperature (22 °C) with a 25 mm diameter stainless steel cone and plate that has a cone angle of 1° and truncation gap of 53 µm. To have a good estimation of the true bulk viscosity, 0.1% polysorbate 80 was added as a surfactant to the solutions. The addition of the surfactant prevented the formation of protein aggregation on the surface of solutions[50].

### 4.5 Droplet Volume and Velocity Measurements

To measure the average volume of individual droplets, 1,000 droplets of various solutions, including thrombin and the solutions mentioned in Section 4.4, were expelled using a micro-valve dispensing device (The Lee Company, cat. no. INKX0517500A) mounted on a custom-designed droplet-based bioprinter (Jetlab® 4, MicroFab Technologies). The mass was measured using a balance (Excellence Plus XP analytical balance, Mettler-Toledo International, Columbus, OH) and the density was calculated by measuring the mass of a 100 L solution in an Eppendorf tube. For velocity measurements, the deposition of droplets was recorded using an in-built camera and analyzed using ImageJ software (National Institutes of Health, Bethesda, MD).

### 4.6 Droplet-based bioprinting for in-vitro Fabrication of Skin Constructs

The custom-designed droplet-based bioprinter was placed in a laminar flow cabinet (Air Science Purair VLF36) for sterility purposes. Three micro-valve dispensing devices (The Lee Company) were used with a removable nozzle (250-m diameter; The Lee Company, cat. no. INZA3100914K), which can be separately controlled to dispense individual solutions. For three fluid reservoirs, fibrinogen with/without adECM were maintained at room temperature and thrombin+Fxiii was kept at 4 °C until deposition. Bioprinting was performed using 5V dwell voltage, 500 s dwell time, 1 s rise/fall time, 0 V echo voltage, 20 s echo time, and 100 Hz frequency. Pneumatic pressure of 126 kPa was applied to flow solutions from the fluid reservoirs. A layer of fibrinogen with various compositions (with/without adECM and ADSCs) and another layer of thrombin with Fxiii (as a crosslinker) was alternatively bioprinted in droplets to build 3D skin constructs. To elaborate, on an 18 × 18 mm glass slide, fibrinogen was deposited and coalesced in a 5-mm circular shape. Then, thrombin was bioprinted on the coalesced structure in the same dimension to crosslink fibrinogen oligomers. The process was repeated three times to achieve the desired thickness.

### 4.7 Scanning Electron Microscopy (SEM) imaging

Morphological characterization of fibrin constructs with and without Fxiii at different concentrations (0, 1, or 2%) was visualized using Scanning Electron Microscopy (SEM) (FEI Nova NanoSEM 630 FESEM, FEI Company, Hillsboro, OR). First, samples were fixed with 4% paraformaldehyde in DPBS overnight at 4 °C. The following day, samples were washed trice with 1× DPBS for 5 min each and then washed thrice for 20 min using HyPureTM molecular biology grade deionized distilled (DDI) water (GE Healthcare Life Sciences). Next, samples were dehydrated using ethanol diluted with DDI water gradually using 50% ethanol for 5 min, 75% ethanol for 5 min, 90% ethanol for 5 min, 95% ethanol for 5 min and 100% ethanol twice for 20 min each. Thereafter, samples were placed inside a critical-point dryer (Leica EM CPD3000, Leica Microsystems, Wetzlar, Germany). Dried samples were then coated with carbon tape using the EM ACE600 sputter coater (Leica Microsystems) and imaged under a high vacuum mode using the Everhart-Thornley detector at an accelerating voltage range from 3.0 to 7.0 keV with a working distance from 3.8 to 6.8 mm.

### 4.8 Cell Viability of ADSC-laden Skin Constructs

LIVE/DEAD staining was carried out using Calcein AM (Thermo Fisher Scientific) and ethidium homodimer-1 (EthD-1, Invitrogen) solutions. Bioprinted constructs were stained on Days 1 and 7 and imaged under a Zeiss Fluorescence Microscope (White Plains, NY) via 2X and 20X magnifications, where dead cells took up the red staining (EthD-1) and live cells took up the green staining (Calcein AM). Images were then taken under Alexa Fluor 568 nm for red fluorophores and Alexa Fluor 488 nm for green fluorophores. Cell viability (%) was determined after the deconvolution process to reduce the background signal based on the number of cells, which had green or red fluorescence by dividing the number of green-fluorescent cells with the total number of cells and multiplied by 100.

### 4.9 Oil Red Staining

To evaluate the intracellular lipid accumulation at different concentrations of adECM (0, 1, or 2%), 1 million/mL of ADSC-laden constructs were bioprinted in well plates and cultured in growth or differentiation media for 14 or 28 days. After the culture, constructs were fixed using 10% neutral buffered formalin for 10 min at room temperature, washed twice with PBS, and stained with 1 mL of 0.5% Oil-red O solution (Sigma-Aldrich, St. Louis, MO) for 15 min, then repeatedly washed with distilled water to remove the background of Oil-red O stain. Representative images were obtained with the EVOS microscope. The area ratio (%) of Oil-red O staining as determined by Image J.

### 4.10 LipidTox Staining of Bilayer Constructs

To assess the intracellular lipid accumulation at different concentrations of adECM (0, 1, or 2%), bilayer constructs were 3D bioprinted, where the first layer was bioprinted with 1 million/mL ADSCs and adECM (0, 1, or 2%) and the second layer was bioprinted on top of the first layer at an ADSC density of 1 million/mL without adECM. The bilayer constructs were cultured in growth media for 14 or 28 days. After culture, constructs were fixed with 4% paraformaldehyde and washed with PBS and stained for LipidTox (Invitrogen 1:500) and DAPI (ThermoFisher, Waltham, MA) using a Zeiss Fluorescence Microscope (White Plains, NY). Mean cellular intensities were quantified using Image J. Lipid droplets were counted positive for adipogenesis if they contained at least one lipid-positive cell.

### 4.11 Surgical Procedures

The surgical procedures were approved by and performed according to guidelines established by American Association for Laboratory Animal Science (AALAS) and Institutional Animal Care and Use Committee (IACUC, protocol #46591). A total of 48 10-12 weeks old female RNU nude rats were obtained from Charles River Laboratories and cared for in our animal facility. We trained the animals in order to have them wearing Elizabethan collars (E-collars) (Kent Scientific, Corporation, Torrington, CT) over ten days before the surgery as required to prevent the animals from accessing their scalps for grooming, which could hamper the integrity of dressing and the reconstructed tissue at the wound site. Rats were gradually trained with E-collars for 30 min on Day 1, to 1 h on Day 2, and eventually 24 h on Day 10 until they fully tolerated E-collars.

Rats were anesthetized with an intraperitoneal injection of ketamine (Midwest Veterinary, Lakeville, MN) mixed with xylazine (LLOYD Inc., Shenandoah, IA) at a dose of 100 mg/kg for ketamine and 10 mg/kg for xylazine, if necessary, after inhalation of isoflurane (2-5%) over 3-5 min. When animals were fully anesthetized, artificial tears (Rugby Laboratories, Corona, CA) were placed on both eyes, and heads were shaved and treated with betadine surgical scrub followed by ethanol. Bupivacaine (0.015 mg/kg) at a concentration of 2.5 mg/mL was injected at the site of incision under the skin (∼0.15 - 0.2 mL) prior to the surgery. Ethiqa XR^TM^ (Fidelis Pharmaceuticals, North Brunswick, NJ) (0.65 mg/kg) was also administered. Two 6-mm (diameter) full-thickness (2 mm depth) skin defects were generated using a 6-mm punch biopsy on the parietal bone. The spacing between the wounds was maintained at ∼3 mm. DBB was utilized to bioprint two compartments including the hypodermis and dermis. The same bioprinting procedures and parameters were used as described for fabricating the in-vitro skin constructs. To elaborate, for the hypodermis compartment, the bioink was deposited with a spacing of 0.8 mm between droplets, and then the crosslinker was deposited with a spacing of 0.4 mm between droplets. This process was repeated thrice. Followed by 15 min of crosslinking at room temperature, the dermis compartment was bioprinted with a spacing of 0.6 mm between droplets and crosslinker was deposited with a spacing of 0.4 mm between droplets. Again, the layers were deposited thrice. Then, the bioprinted constructs were allowed to crosslink for an additional 10 min at room temperature. After the surgery, Tegaderm^TM^ transparent dressings (3M Company, St. Paul, MN) were used to cover the bioprinted sites without suturing, and E-collars were put around the necks of animals to prevent them from accessing wound sites. The rats were able to function normally after the procedure. Animals were observed and weighed daily for at least 10 days post-surgery. Animals were briefly anesthetized with isoflurane daily to allow imaging of the defect sites for 10 days and every other day beyond Day 10. Tegaderm and E-collars were removed after 7 and 14 days, respectively. Animals were euthanized at Week 2 or 4 for tissue histology and immunohistochemistry to evaluate skin regeneration.

### 4.12 Morphometric Evaluation of Skin Re-epithelialization and Closure

Wound healing was characterized by taking photographic images of the wounds every two days for four weeks. The images were then analyzed using ImageJ Fiji. Briefly, a freehand tool of the program was used for mapping the contours of wounds, and the wound area was measured at each predetermined time-points. From the measured areas, the percentages of wound closure (%) and skin re-epithelization (%) were calculated according to a previously published study[30] using (Equations 1) and (2);

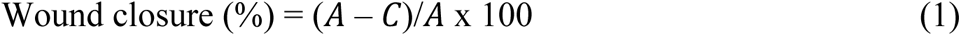

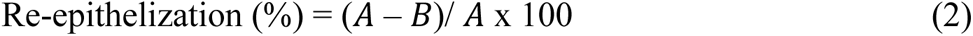

where the wound area at the time of surgery was defined as “*A*” and the unclosed wound area at a specific time-point was defined as “*C*”. Further, the non-re-epithelialized but thinner skin area at a specific time-point was defined as “*B*”. The percentage (%) of re-epithelialization and percentage (%) of wound closure for each group were plotted as a function of time (days) using Prism 8 software (GraphPad).

### 4.13 Histological Assessment

The excised wound and surroundings (∼ 6 mm) were fixed in 4% formaldehyde for 48 h, embedded in O.C.T cryomatrix (Thermo Fisher Scientific) and sequentially sectioned using a Leica CM1950 cryostat (Leica Biosystems) at −32 °C with a section thickness of 8-10 µm. First, the sectioned samples were placed on a Hematoxylin and Eosin (H&E) automated staining platform (Leica Auto Stainer XL, Leica Biosystems) without applying any heat during the staining process. Using Neo-Mount® anhydrous mounting medium, H&E stained sections were mounted and imaged using a Keyence BZ-9000 fluorescence microscope (Keyence Corporation of America, Elmwood Park, NJ) under brightfield using a neutral density optical filter. Based on the H&E stained images, the number of downgrowth of hair follicles in the epithelial layer was counted. To evaluate the deposition of collagen histomorphometrically, sectioned samples were stained using Masson’s Trichrome staining (MTS) kit (Sigma-Aldrich, cat. No. HT15-1KT) based on the manufacturer’s protocols. The sectioned samples were dehydrated according to the protocol and then mounted using Neo-Mount® anhydrous mounting medium (Millipore) and imaged by the Keyence BZ-9000 fluorescence microscope as described before. Using the images of MTS, collagen deposition was quantified. The collagen index was determined according to Eq. (3) from the literature [51],

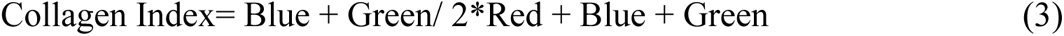

where, Blue, Green, and Red indicate pixel intensities. Images were processed using Image J and pixel intensities were determined using the histogram of Red Green Blue (RGB) color images, where the mean intensity of blue, green, and red pixels was used in calculations. The collagen index was determined to be between 0 to 1, where ‘1’ indicates complete Blue and ‘0’ indicates complete Red. The epidermis layer was not included in measurements. A total of one image per each stained section and a total of three stained sections from each wound area were used for measurements for each group.

### 4.14 Immunohistochemistry (IHC)

IHC staining was conducted after thawing sectioned samples for 10 to 20 min at room temperature from −30 °C. To investigate adipogenesis in the hypodermis compartment, High-Content Screening (HCS) LipidTox Red neutral lipid stain (Invitrogen, cat. No. H34476, 1:500) against neutral lipid droplets marker was used. All cryo-sectioned samples were treated with blocking solution of 10% (w/v) normal goat serum (NGS) (diluted in distilled water) to avoid non-specific binding sites. Thereafter, LipidTox and DAPI (1:500) were treated for 30 min at room temperature in the dark. After that, a mounting medium was applied to the sectioned samples. Fluorescence images were obtained using the Zeiss Fluorescence Microscope as described before with 20× magnification. Mean cellular intensities were quantified using Image J.

To investigate vascularization in sections, cluster of differentiation 31 (CD31) antibody against the murine endothelial cell surface marker was used. All cryo-sectioned samples were treated with a blocking solution of 10% (w/v) NGS (diluted in distilled water) to avoid non-specific binding sites. Thereafter, the primary anti-CD31 antibody (Abcam, cat. No. ab28364, 1:200) was treated overnight at 4 °C in the dark. After washing several times with PBS, the secondary antibody conjugated to Alexa Fluor® 647 goat anti-rabbit IgG (H+L) (Invitrogen, cat. No. A-21245, 1:500) was then treated for 1 h to visualize the antigen. Finally, DAPI mounting medium was applied to sectioned samples to visualize cell nuclei and stored in the dark at 4 °C. Fluorescence images were obtained using the Zeiss Fluorescence Microscope with 20× magnification. Mean cellular intensities were quantified using Image J.

To confirm the identity of the large adipocyte-like structures in the wound bed, staining for anti-murine Fatty acid binding protein 4 (Fabp4) (R&D Systems, AF1443) was used. This was performed in parallel with staining for GFP (Abcam 13970) to confirm the presence of ADSCs in the wound bed. All tissue samples were stained with the horizontal whole mount method as we described before[52]. Full thickness skin samples were sectioned and fixed in 4% paraformaldehyde, frozen in OCT, and sectioned at 60 m. Sections underwent blocking with gelatin blocking buffer, followed by staining with primary antibodies overnight at 4 °C and secondary antibodies for 4 h. Samples were then mounted onto a glass slide with glycerol. Imaging was performed at 20x with the SP8 Confocal Microscope (Leica, TCS SP8 X) at the WSU Franceschi Microscopy and Imaging Center (FMIC).

### 4.15 Statistical Analysis

All data were presented as mean ± standard deviation unless stated otherwise. To compare three or more groups, data were analyzed by one-way ANOVA with post hoc Fisher’s individual tests for differences of means at a 95% confidence level. In order to evaluate % re-epithelialization and wound closure, one-way ANOVA with post hoc Fisher’s individual tests (assumption of equal variances) was used at a 95% confidence level in order to determine significance levels based on the area under the curve for each group from Day 0 to 28. Differences were considered significant at p* < 0.05, p** < 0.01, and p*** < 0.001. All statistical analysis was performed using Prism 9 (GraphPad).

