## Supporting Information for "Intraoperative Bioprinting of Human Adipose-derived Stem cells and Extra-cellular Matrix Induces Hair Follicle-Like Downgrowths and Adipose Tissue Formation during Full-thickness Craniomaxillofacial Skin Reconstruction"

### Supplementary Figures

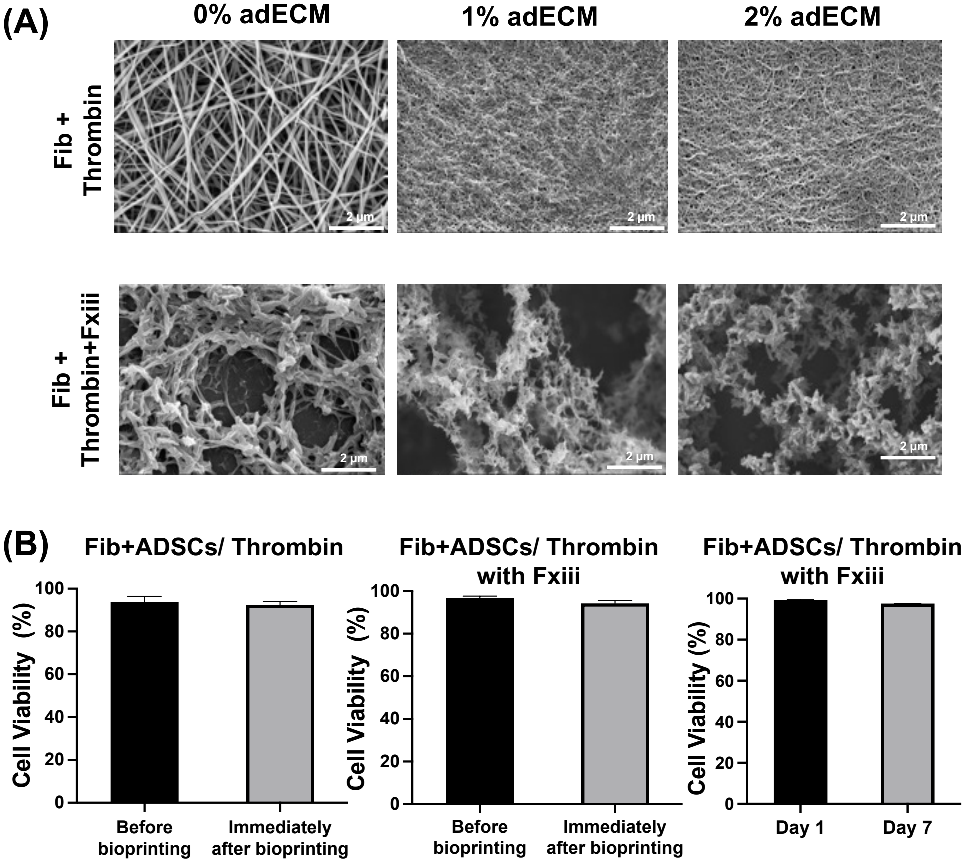


**Figure S1.** (A) SEM images of fibrinogen containing 0, 1 or 2% adECM crosslinked with thrombin in the presence and absence of Fxiii. (B) Viability of ADSCs-laden in fibrinogen (at a density of 1 million/mL) in the absence (left) and presence (middle) of Fxiii before and after bioprinting and in the presence of Fxiii (right) at Days 1 and 7 (*n*=3, 'Fib' indicates fibrinogen).

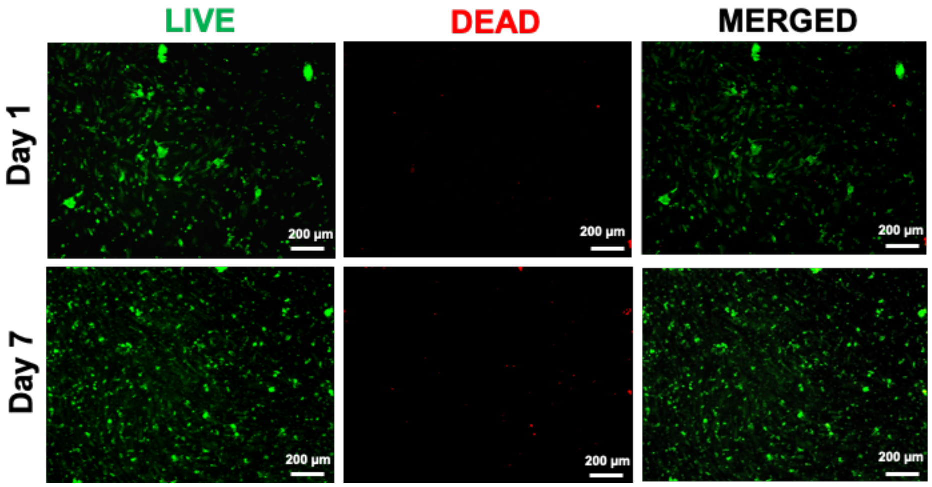


**Figure S2.** LIVE/DEAD images of ADSCs-laden fibrin (1 million/mL cell density) at Days 1 and 7 after bioprinting (*n*=3).

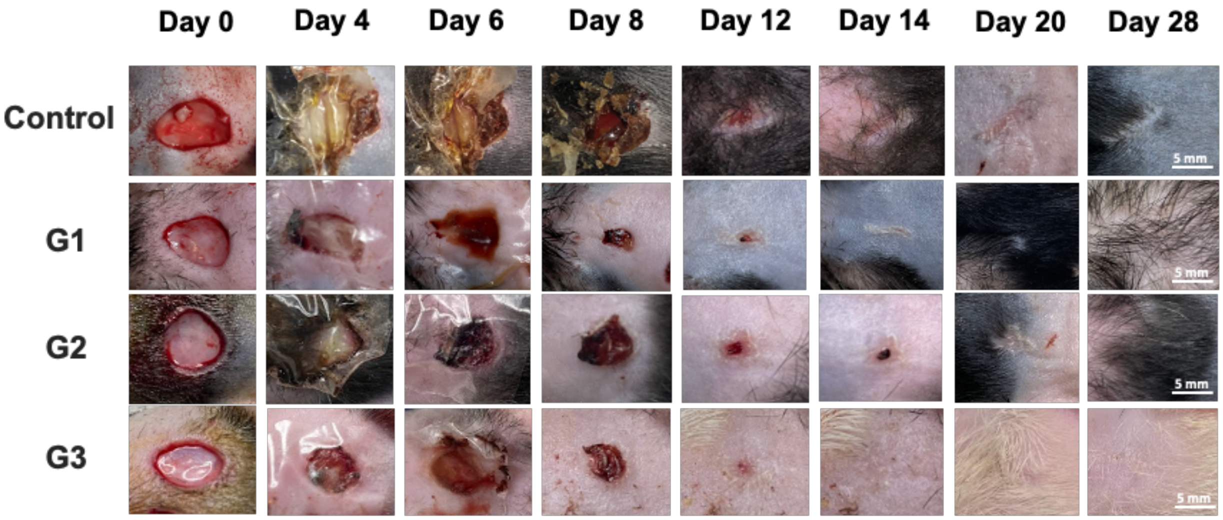


**Figure S3.** Representative wound images, from Day 0 to 28 for Study #1.

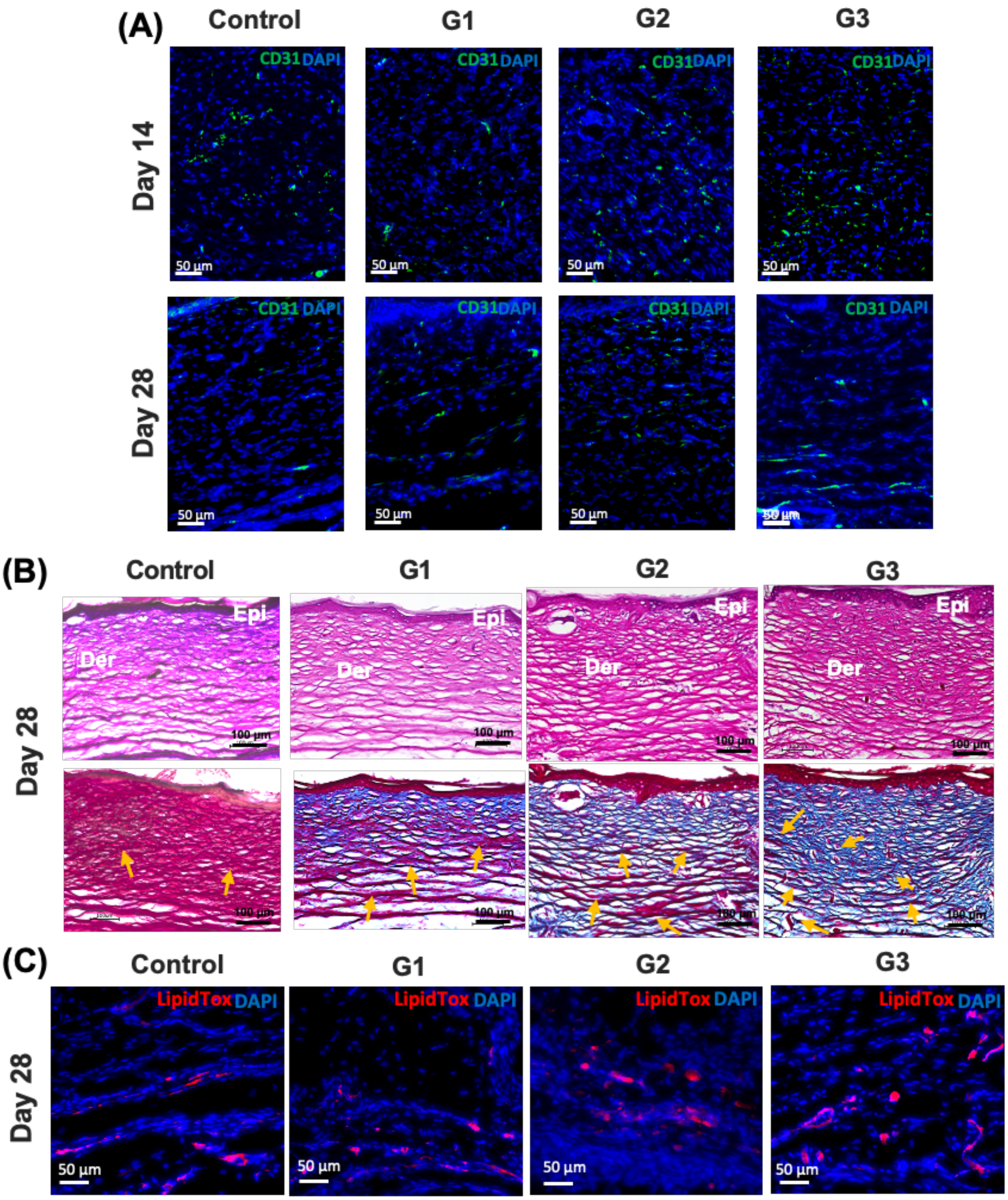


**Figure S4.** (A) CD31 immunostaining images at Days 14 and 28 for Study #1. (B) H&E and M&T staining images at Day 28 for Study #1. (C) LipidTox staining images at Day 28 for Study #1. Yellow arrows indicate blood vessels.

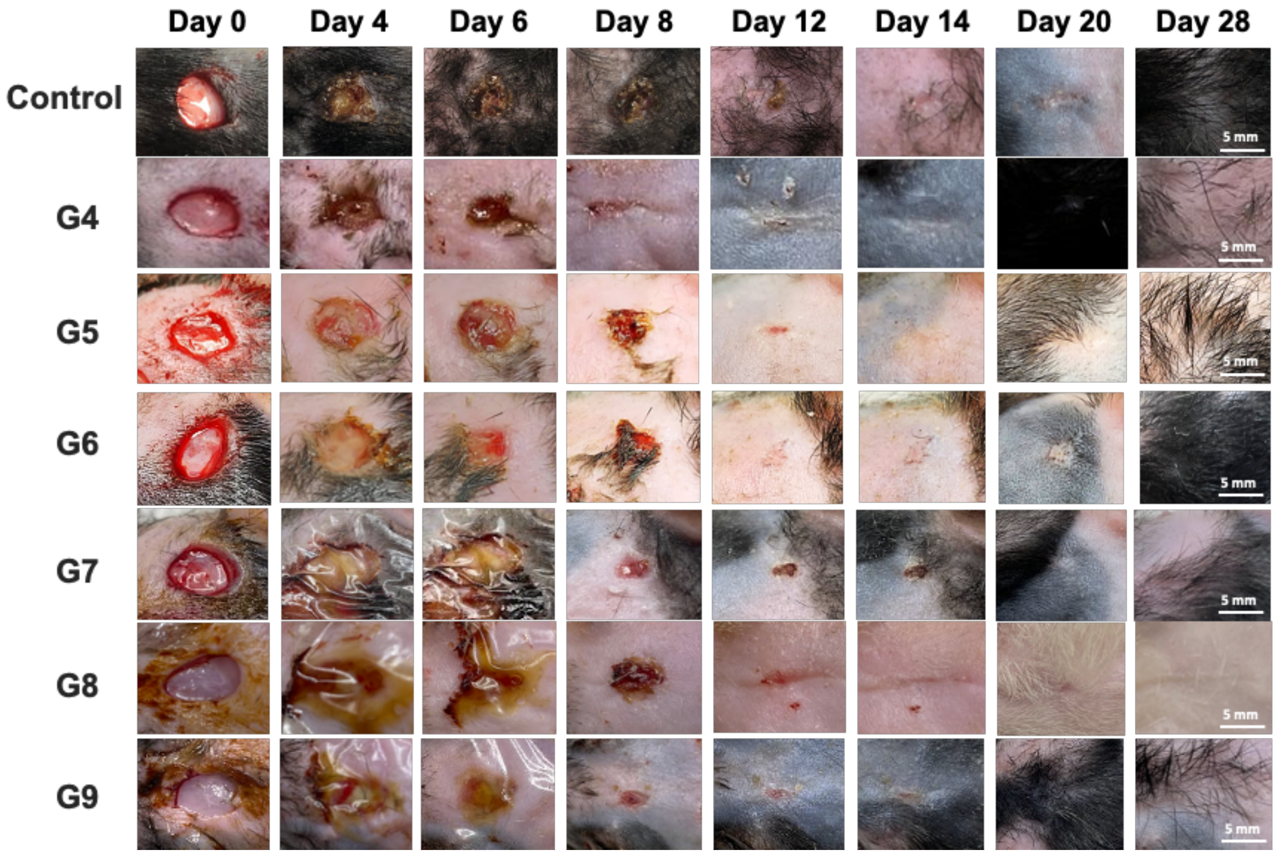


**Figure S5.** Representative wound images, from Day 0 to 28 for Study #2.

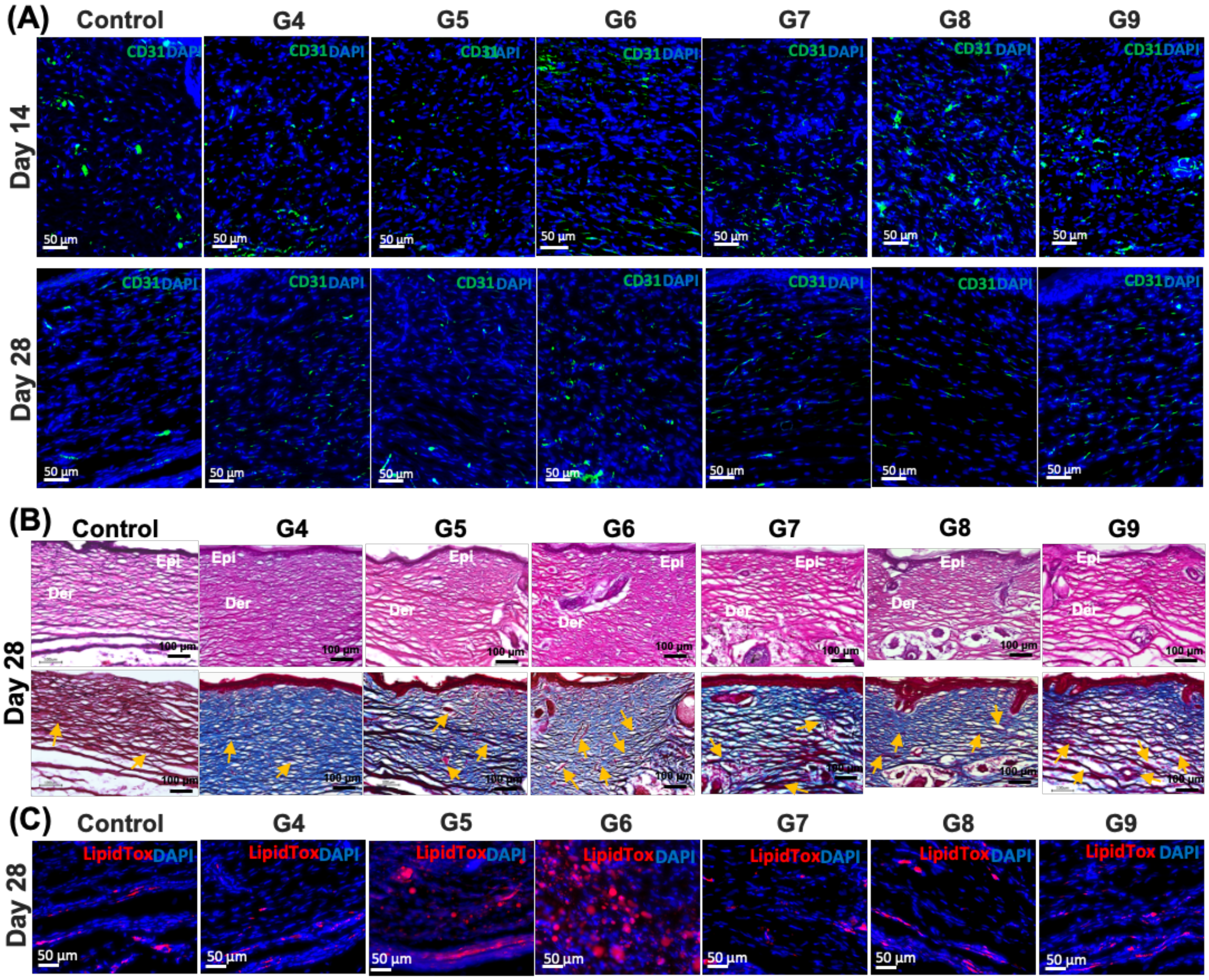


**Figure S6.** (A) CD31 immunostaining images at Days 14 and 28. (B) H&E and M&T staining images at Day 28 for Study #2. (C) LipidTox staining images at Day 28 for Study #2. Yellow arrows indicate blood vessels.

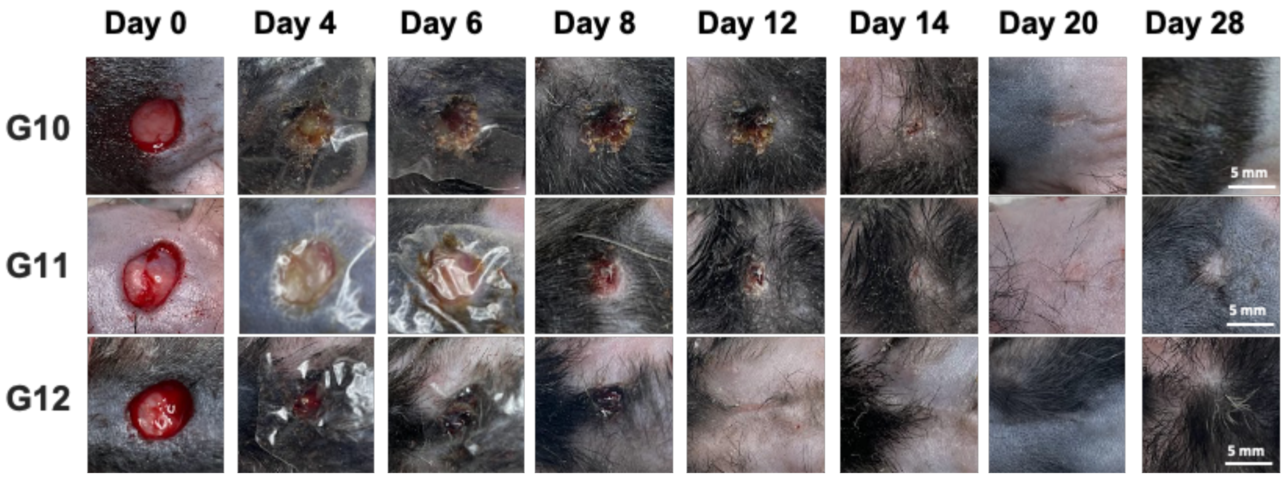


**Figure S7.** Representative wound images, from Day 0 to 28 for Study #3.

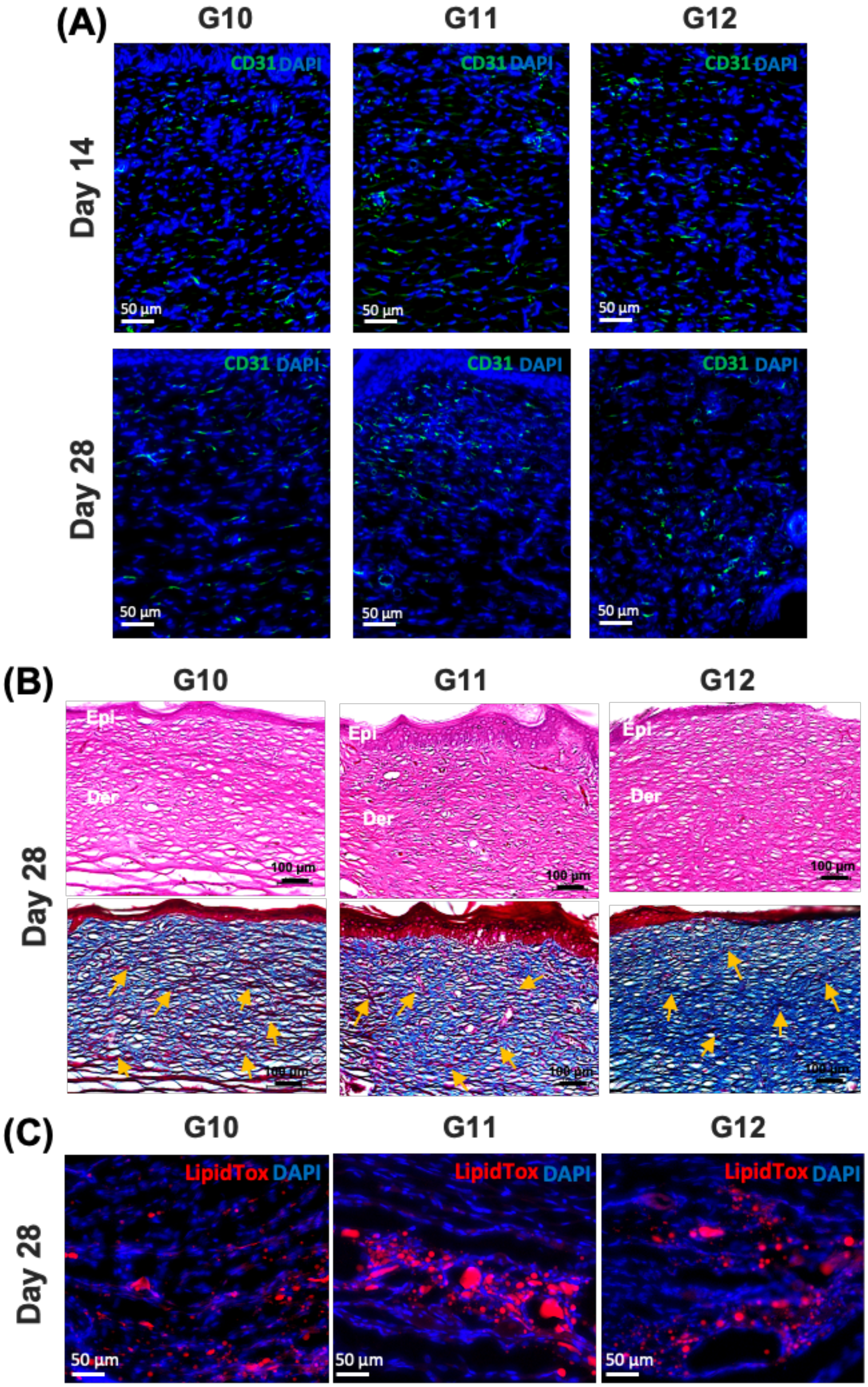


**Figure S8.** (A) CD31 immunostaining images at Days 14 and 28. (B) H&E and M&T staining images at Day 28 for Study #3. (C) LipidTox staining images at Day 28 for Study #3. Yellow arrows indicate blood vessels.

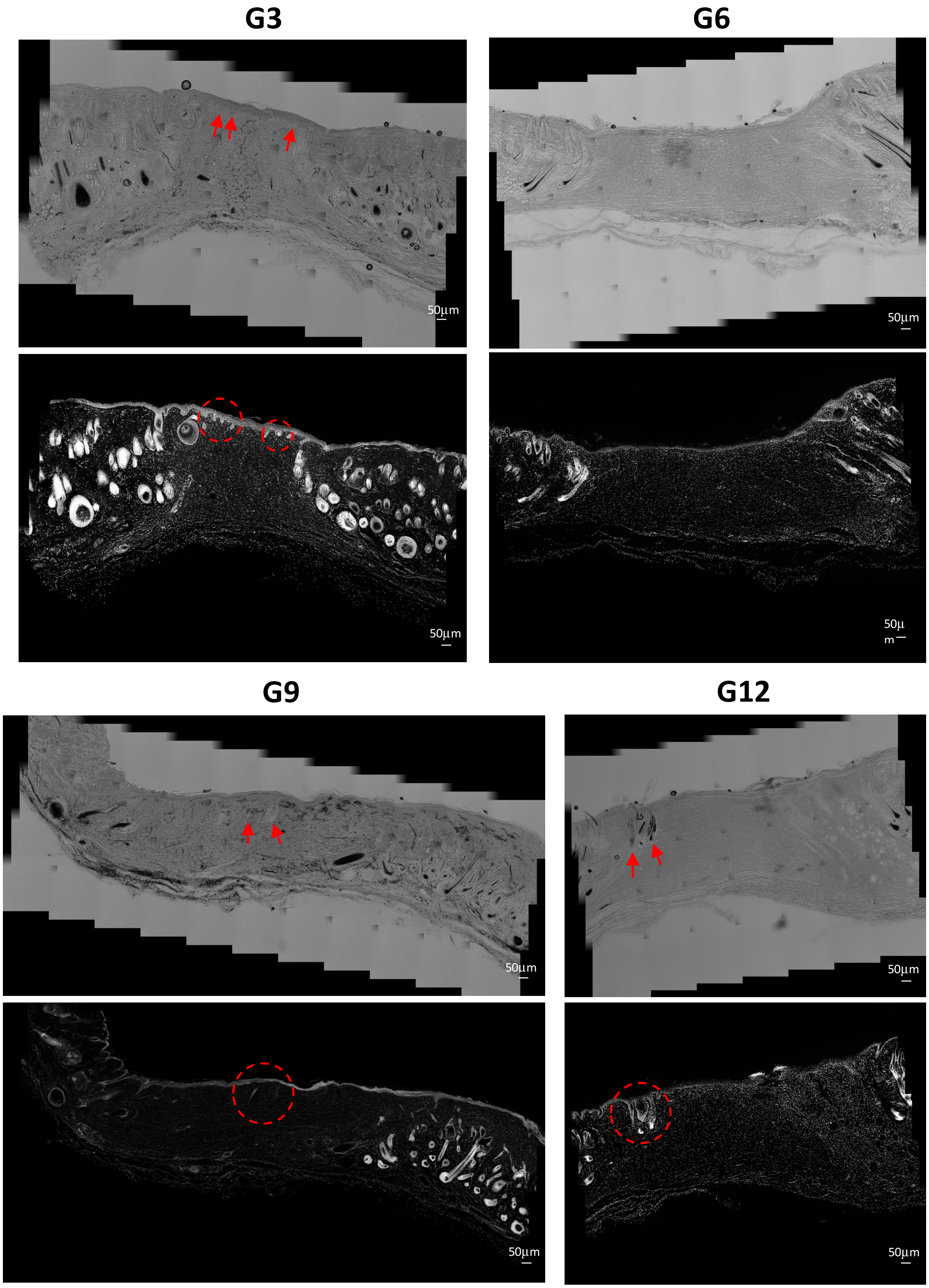


**Figure S9.** Optical images (up) and dark field images (down) of the reconstructed skin in G3, G6, G9 and G12 at Day 14. Red arrows indicate the downgrowth of hair follicles.

### Supplementary Videos

**Video S1**: IOB of the skin reconstruction for G3 in real time.

**Video S2**: IOB of the skin reconstruction for G6 in real time.

**Video S3**: IOB of the skin reconstruction for G9 in real time.

**Video S4**: IOB of the skin reconstruction for G12 in real time.
